# LIPA-driven hydrolysis of cholesteryl arachidonate promotes cancer metastasis via NF-κB

**DOI:** 10.1101/2022.03.11.484042

**Authors:** Zhicong Chen, Fukai Chen, Hyeon Jeong Lee, Meng Zhang, Xianglin Yin, Mingji Dai, Ji-Xin Cheng

## Abstract

Cholesteryl ester (CE) is an established marker in many types of aggressive cancers. Yet, the function of CE homeostasis during cancer progression is largely unknown. Here, enabled by Raman spectromicroscopy, pan-cancer bioinformatic analysis, and biocompatible Raman probe of cholesterol, LIPA was identified as the key regulator of CE hydrolysis in cancer cells. LIPA inhibition induced aberrant accumulation of CE-rich lipid droplet at single cell level. LIPA-driven CE hydrolysis was directly visualized by stimulated Raman scattering imaging of alkyne-tagged cholesterol. Cholesteryl arachidonate was identified as a dominant substrate that is rapidly hydrolyzed by LIPA. Inhibition of LIPA effectively suppressed cancer metastasis both in vitro and in vivo. Mechanistically, LIPA inhibition suppresses NF-κB signaling, while the NF-κB members positively regulate the expression level of LIPA, indicating regulation of CE homeostasis by a LIPA-CE-NFκB feedback loop. Collectively, our findings reveal that LIPA drives CE hydrolysis during cancer progression and is an important metabolic target for cancer therapy.

## Introduction

Aberrant cholesterol metabolism is a vital hallmark of cancer, including increased cholesterol biosynthesis, enhanced uptake, and derivatives enrichment^1,2^. Cholesteryl ester (CE), as an important form of cholesterol storage and transportation, is redundantly accumulated in a variety of tumors, which contributes to tumor progression and poor prognosis^3–5^. CE-rich lipid droplets (LDs) function as a reservoir for neutral lipids, not only to reduce the toxicity of excessive cholesterol, but also to hydrolyze under increased demand. It is well defined that cholesterol is esterified into CE by SOAT1 in tumor, and PTEN-PI3K-Akt^3^, Wnt–β-catenin^6^ and SREBP signaling^5^ have been implicated in SOAT1-related cancer progression. Naturally, SOAT1 has thus been highlighted as a metabolic target of cancer^7^. However, CE hydrolysis in cancer is under studied in comparison with cholesterol esterification. Consequently, the functional role of CE in cancer development remain elusive.

CE hydrolysis, catalyzed by lipases, ensures rapid obtainment of free cholesterol and fatty acids which are required in cancer development^2,8^. Learning from the studies of atherosclerosis in which CE accumulation is a typical hallmark^12^, neutral lipolysis of CEs are mediated by CES1, NCEH1 and LIPE^9,10^, while LIPA catalyzes acidic lipolysis in lysosomes^10,11^. Compared with the extensive studies in atherosclerosis^12^, there are only limited studies on CE lipases in cancer. NCEH1 was correlated with cancer cell invasiveness^13^. Increased LIPA expression was found in renal cancer and was shown to promote cell proliferation and survival via activation of the Src/Akt pathway^14^. CES1 overexpression exerted an anti-proliferative effect in liver cancer cells^15^. These studies point to the functional alternation of CE lipases in cancer. Despite these initial efforts, the dominant substrate of hydrolysis and the regulatory mechanism of CE hydrolysis in cancer progression remain unknown.

Here, we identified LIPA and NCEH1 as potential regulators of CE hydrolysis during cancer development using the database of cancer patients. The expression of CE lipases was correlated with SOAT1 and prognosis. The robust hydrolysis activity of LIPA was characterized by stimulated Raman scattering (SRS) imaging and confocal Raman microspectroscopy measurement of intracellular CE. Inhibition of LIPA induced accumulation of CE-rich LDs. Importantly, we tracked the LIPA-driven CE hydrolysis process directly by bio-orthogonal chemical imaging using alkyne-tagged cholesterol^16^, verifying LIPA as the major CE lipase in cancer. Furthermore, cholesteryl arachidonate was identified as the major CE species hydrolyzed by LIPA using mass spectrometry. Suppression of LIPA significantly reduced cancer metastasis both in vitro and in vivo. Mechanistically, LIPA-CE-NFκB feedback loop was found to promote cancer metastasis. Overall, our findings put forward LIPA as the vital lipase for cholesteryl arachidonate during cancer metastasis, revealing a new metabolic target for cancer treatment.

## Results

### Genetic assay identifies regulators of CE hydrolysis during cancer progression

To identify the major regulator of CE hydrolysis, cancer cohorts in The Cancer Genome Atlas (TCGA) were interrogated. CE homeostasis results from the balance between esterification and hydrolysis. CE hydrolysis is a reverse reaction of CE esterification manipulated by SOAT1. We thus studied the correlation between SOAT1 and several CE lipase candidate genes (NCEH1, LIPA, LIPE, CEL, and CES1) in cancer. Among them, LIPE, CEL, and CES1 expression levels showed weak correlations with SOAT1, while SOAT1 showed strong positive correlations with NCEH1 and LIPA, especially with LIPA (**Fig. 1a-b**). Significantly positive correlations between SOAT1 and NCEH1/LIPA were found in almost all cancer cohorts from TCGA. Then, the gene expression levels were further characterized in different comparisons, including tissue subtype and patient prognosis. Consistent with previous studies^2^, SOAT1 expression is upregulated in cancer tissues in comparison with normal/paracancerous tissues and contributes to unfavorable overall survival in multiple cancer types (**Fig. 1c**). Similarly, the tumor-promoting expression patterns were also found in NCEH1 and LIPA. Increased NCEH1/LIPA expression in cancer contributes to poorer prognosis (**Fig. 1d-g**). Meanwhile, the other candidate genes were generally downregulated and showed inconsistent prognosis effects (**Fig. 1c**). Collectively, the strong positive correlations with SOAT1 and cancer-promoting expression pattern suggest that NCEH1 and LIPA could serve as regulators of CE hydrolysis during cancer progression.

**Fig. 1.**
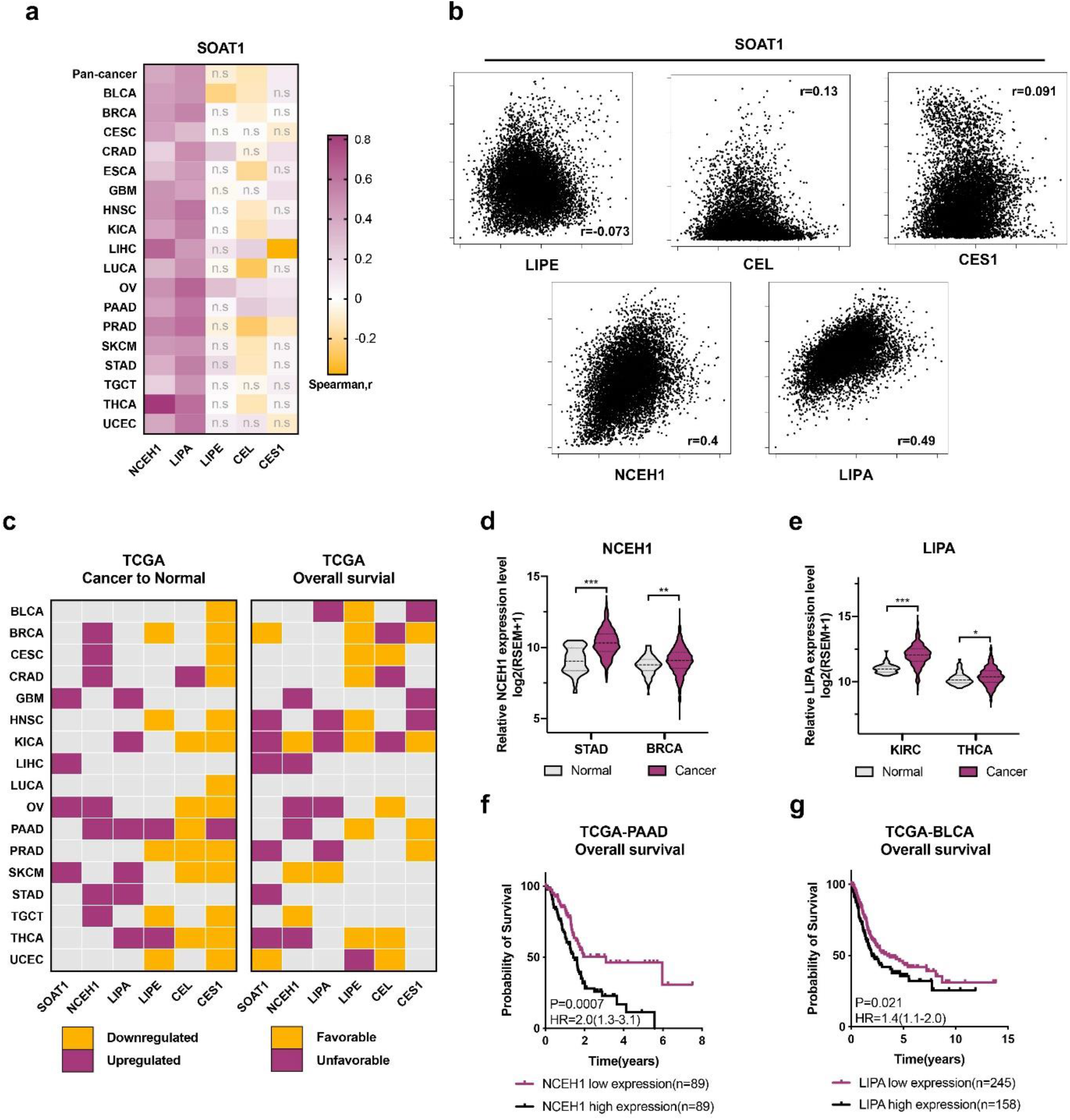
CE hydrolysis contributes to cancer progression. **a** A Heatmap of Spearman’s rank correlation coefficient between expression level of SOAT1 and CE lipases in different cancer cohorts from TCGA. **b** Correlation plot between SOAT1 and CE lipases in pan-cancer cohorts of TCGA. **c** The expression microarrays of SOAT1 and CE lipases in tissue type and prognosis effect. **d-e** CE lipases expression between normal and cancer tissues in representative cancer cohorts. NCEH1(**d**) and LIPA (**e**). **f-g** Kaplan-Meier curves of the CE lipases expression level on OS in representative cancer cohorts. NCEH1(**f**) and LIPA (**g**).

### LIPA is the functional regulator of CE hydrolysis in cancer cells

CEs are stored in LDs. Increased LD formation is a typical and immediate signature for aberrant CE accumulation resulting from either the redundant CE esterification or the inhibition of CE hydrolysis. To verify the CE-hydrolysis ability of LIPA and NCEH1, their selective inhibitors (Lalistat 1-LIPA, JW480-NCEH1)^17,18^ were adopted. A high-speed femtosecond stimulated Raman scattering (fs-SRS) microscope (Supplementary **Fig. 1**) was deployed to image and quantify the amount of LDs at single cell level. By tuning the laser beating frequency to be resonant with C-H stretching vibration, stimulated Raman loss (SRL) signals arose from C-H rich biomolecules (**Fig. 2a**), such as intracellular LDs^19^. To identify the common CE hydrolase among different cancers, totally four types of cancer cells were studied, including T24 bladder cancer, MDA-MB231 breast cancer, MiaPca2 pancreatic cancer, and PC3 prostate cancer. Except for T24, the others have been highlighted with a high amount of CE in previous studies^3,4,20^. Due to the great potential of LIPA in CE hydrolysis, Lalistat 1 was first applied to suppress CE hydrolysis. LDs in all four types of cancer cells dramatically increased and were largely dispersed around the endoplasmic reticulum after treatment with Lalistat 1. Furthermore, such enrichment of LDs can be restored by recovering cells from Lalistat 1 in inhibitor-free media (**Fig. 2b-c**). Among four cells, the fold change of both LDs accumulation and recovery in T24 was most obvious. The LD hydrolysis rate was quantitated (**Fig. 2d**) by using intensity ratio of Lalistat 1 treatment to 24 h recovery (10.83 in T24, 3.45 in MDA-MB231, 2.20 in MiaPca2 and 1.51 in PC3). Collectively, Lalistat 1 could effectively induce the reversible accumulation of intracellular LDs.

**Fig. 2.**
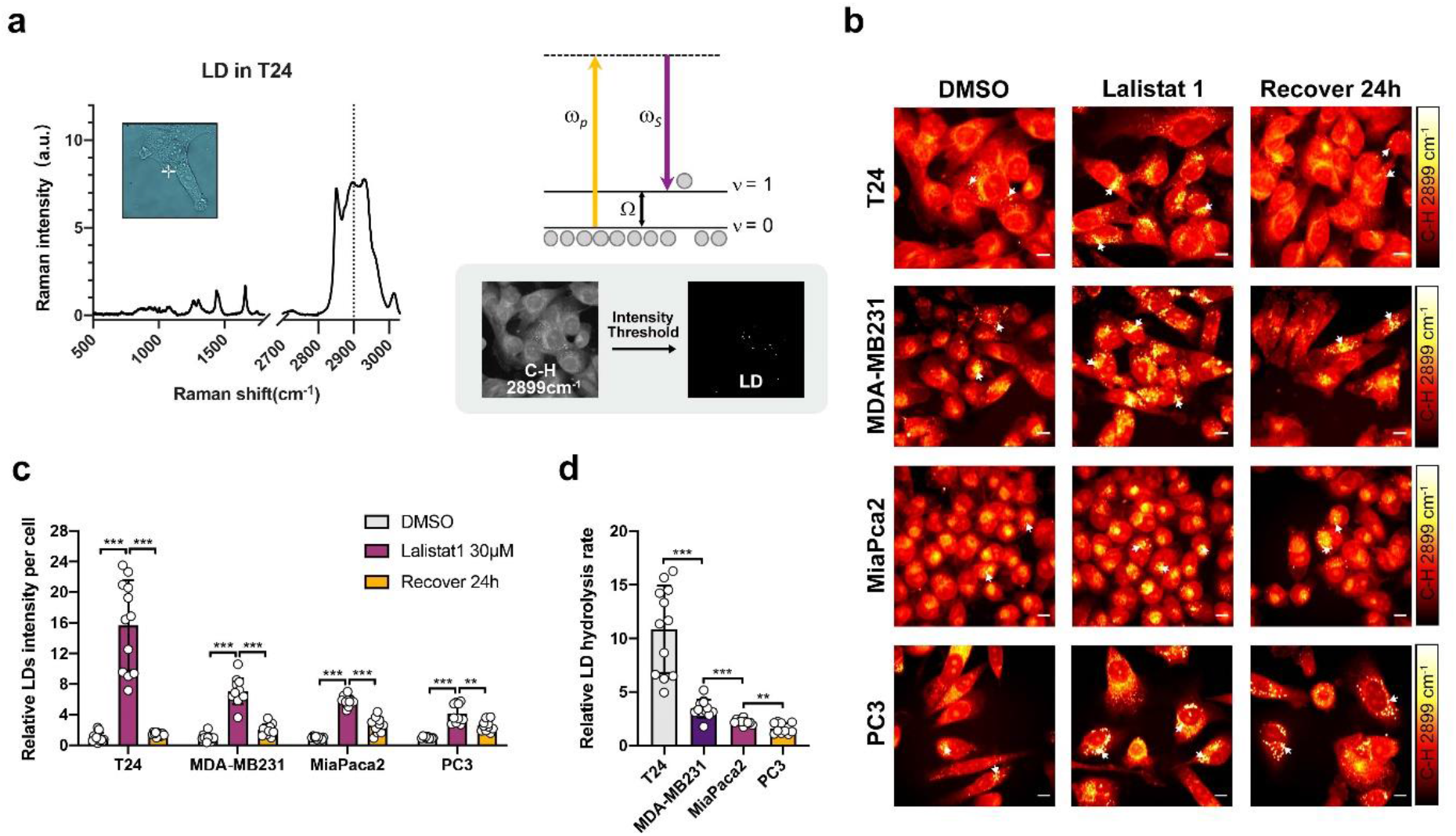
LIPA inhibition by Lalistat 1 induces the reversible accumulation of intracellular lipid droplets. **a** Schematic illustration of LD detection by fs-SRS. High Raman intensity of LD was detected in C-H vibration region. SRS is a dissipative process in which energy corresponding to the beating frequency (ω_p_ – ω_S_) is transferred from input photons to a Raman-active molecular vibration (Ω). LDs were further picked up by applying a threshold out of total cellular area. **b-c** Representative SRS imagines of four cancer cells after Lalistat 1 treatment and recovery(**b**). Scale bars, 10μm. Average LDs SRS intensity in each cell (**c**). Each dot represents a single detection frame. At least 100 cells / sample were imaged. **d** Relative LD hydrolysis rate quantitated by using SRS intensity ratio of Lalistat 1 to 24 h recovery. Error bars represent SD of the mean. ** p < 0.01; *** p < 0.001.

Generally, the core of most LDs is enriched with triacylglycerol (TAG) and sterol ester^21^, and is rich in CE in aggressive cancer cells^22^. LIPA functions in the lysosome to catalyze the hydrolysis of CE and TAG^11,14,23^. We next evaluate the preference of LIPA by detecting the content inside the accumulated LDs. Since the dominant fatty acid species of CE and TAG in cancer were both oleate^3,22,24^, the reference spectra of concentration gradient emulsion mixtures with cholesteryl oleate (CE 18:1) and glyceryl trioleate (TAG 18:1) were recorded using confocal Raman spectroscopy. Consistent with previous studies^25^, several characteristic Raman bands of CE were confirmed, including cholesterol rings at 702 cm^-1^, acyl C=C stretching at 1,654 cm^-1^, sterol C=C stretching at 1,670 cm^-1^, sterol CH2 symmetric stretching at 2,870 cm^-1^ and acyl =CH stretching at 3,008 cm^-1^ (**Fig. 3a**). Compared to control cells, all CE signatures were also found in the LDs induced by Lalistat 1, particularly the cholesterol rings at 702 cm^-1^ (**Fig. 3b**). After normalization by the CH2 scissoring and bending band at 1,445 cm^-1^, we generated a calibration curve for CE percentage using the unique cholesterol band at 702 cm^-1^. The peak intensity was linearly proportional to the molar percentage of CE in the emulsions (Supplementary **Fig. 2a**). Based on the calibration, the CE contents of the LDs significantly increased in all cancer cells after Lalistat 1 treatment (**Fig. 3c**), specifically, MDA-MB231 from 13% to 39%, MiaPca2 from15% to 56%, PC3 from 50% to 71%, especially in T24 (from 15% to 70%). Due to the prominent reduction of C=C stretching at 1,654 cm^-1^, we also generated a calibration curve using fatty acids containing different numbers of C=C bonds to quantify the unsaturation degree (Supplementary **Fig. 2b**). The number of C=C bonds in the fatty acids linearly correlated with the normalized peak intensity of C=C stretching vibration at 1,654 cm^-1^ (Supplementary **Fig. 2c**). Lalistat 1 diminished the unsaturation degree of LDs in all four types of cancer cells (**Fig. 3d**). Together, inhibition of LIPA by Lalistat 1 significantly increased the CE percentage and decreased the unsaturation degree in LDs (**Fig. 3e**). Similarly, fs-SRS imaging and Raman microspectroscopy were also performed under NCEH1 inhibition by JW480. JW480 induced LDs accumulation in T24 (**Fig. S3a-b**). However, there was no significant alteration in the CE contents comparing with the control (Supplementary **Fig. 3c-d**), indicating that NCEH1 is not the functional regulator of CE hydrolysis inside cancer cells.

**Fig. 3.**
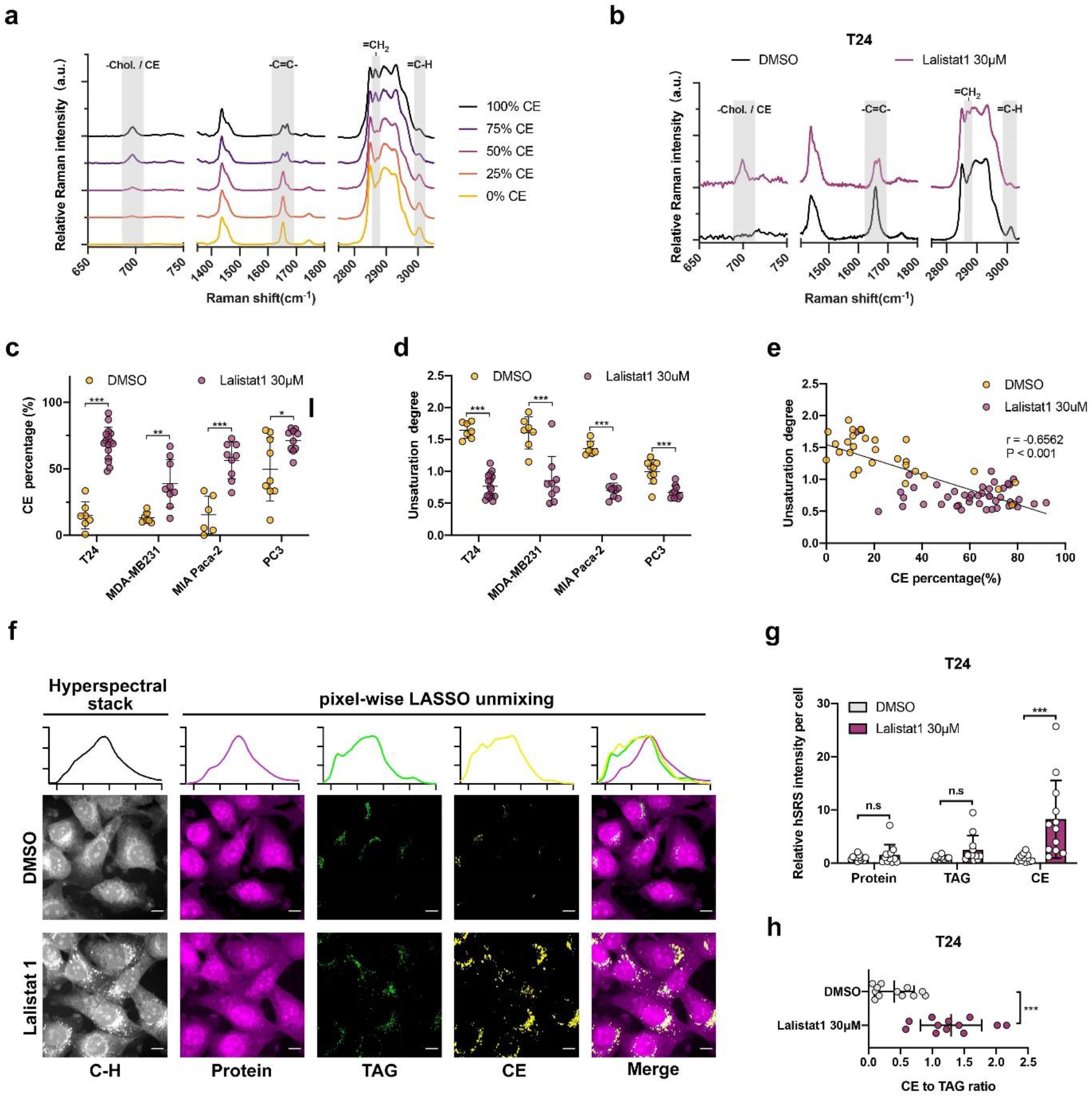
LIPA inhibition effectively impairs CE hydrolysis. **a** Average spontaneous Raman spectra taken from concentration gradient emulsion mixtures with CE 18:1 and TAG 18:1. **b** Average spontaneous Raman spectra taken from LDs in T24 with Lalistat 1 treatment. **c-d** CE percentage (**c**) and unsaturation degree (**d**) in LD of four cancer cells with Lalistat 1 treatment. **e** Correlation between CE percentage and unsaturation degree in LD with Lalistat 1 treatment. Each dot represents a single cell. At least three LDs / cell were detected. **f-g** Representative hSRS imagines and spectra of T24 cells with Lalistat 1 treatment (**f**). Scale bars, 10μm. Quantitative analysis of chemical mapping of T24 with Lalistat 1 treatment (**g**). **h** Quantitative CE to TAG hSRS intensity ratio of T24 with Lalistat 1 treatment. Each dot represents a single detection frame. At least 100 cells / sample were imaged. Error bars represent SD of the mean. * p < 0.05; ** p < 0.01; *** p < 0.001; n.s, not significant.

For high throughput validation in situ, hyperspectral stimulated Raman scattering (hSRS) microscopy was sequentially performed. Bovine serum albumin (BSA), glyceryl trioleate (TAG 18:1), and cholesteryl oleate (CE 18:1) served as the spectral references of protein, TAG, and CE. By tuning the Raman shift frame by frame, 100 images of pure chemical, Lalistat 1-treated, and control T24 cells were separately recorded at the C-H stretching vibration region from 2,800 to 3,050 cm^-1^. Consistent with the fs-SRS imaging results, the abundant LDs in Lalistat 1-treated T24 cells were verified by hSRS. On basis of spectral references, the C-H hyperspectral stacks were decomposed into chemical maps of protein, TAG, and CE by applying pixel-wise LASSO unmixing analysis^26^. For the protein channel, the signals are distributed all over the cell with an enhanced contrast for protein-rich organelles, such as the endoplasmic reticulum and nucleolus. As expected, the signals from both TAG and CE were dominated by LDs distributed around the endoplasmic reticulum (**Fig. 3f**). Compared with the control group, Lalistat 1 boosted up the CE signals robustly, while no significant alteration in protein and TAG is found (**Fig. 3f-g**). The average intracellular CE to TAG ratio increased from 0.4 to 1.3 after Lalistat 1 treatment, demonstrating a switch from TAG-rich to CE-rich phenotype (**Fig. 3h**). Collectively, LIPA inhibition effectively impairs CE hydrolysis, showing that LIPA is the functional CE hydrolysis regulator in cancer cells.

### Visualization of CE homeostasis by SRS imaging of alkyne-tagged cholesterol

To further validate the function of LIPA in CE hydrolysis, we performed fs-SRS imaging of Phenyl-Diyne cholesterol (PhDY-Chol) (**Fig. 4a**). PhDY-Chol is a bio-compatible analog of cholesterol tagged with an alkyne bond (C≡C) that gives a characteristic and strong peak at 2,254 cm^-1^ in the Raman-silent region^16^. The strong signal given by PhDY-Chol has enabled bond-selective imaging of cholesterol esterification and storage inside living cells^16^. Here, the balance between cholesterol esterification and hydrolysis was directly visualized by tracking this analog. Various concentrations of PhDY-Chol (2.5, 5, 10, 20 μM) were applied to monitor intracellular cholesterol trafficking. The central beating frequency between pump and Stokes beams was tuned to be resonant with C-H stretching vibration at 2,899 cm^-1^ and C≡C vibration at 2,254 cm^-1^, where SRL signals arose from C-H rich biomolecules and PhDY-Chol, respectively. LDs that contained PhDY-Chol esters were visible in the C≡C window (**Fig. 4b**). Same as cholesterol, the redundant PhDY-Chol were esterified and stored into LDs to minimize its cytotoxicity (**Fig. 4b-d**). Due to the competitive cholesterol in the complete media, weaker signals were observed in the 2.5 μM group, and the cellular intensity of PhDY-Chol was linearly correlated with its concentration (**Fig. 4e**). After the inhibition of LIPA by Lalistat 1, similar accumulations of LDs were found in C-H windows. At the same time, significantly increased PhDY-Chol LDs were found compared to the control groups regardless of initial PhDY-Chol concentration. The extensive accumulation of PhDY-Chol validates the dominant role of LIPA in CE hydrolysis, providing visual evidence for the tilting of CE balance towards esterification (**Fig. 4c, e-f**). Recovering cells from Lalistat 1 using inhibitor-free media, the aberrant accumulated LDs in both C-H and C≡C windows dramatically reduced, indicating the corrective reboot of CE hydrolysis (**Fig. 4d-f**). Of note, the CE hydrolysis rates were consistent among various concentration groups with the declined PhDY CE percentages to 30% (**Fig. 4g**), suggesting steady CE hydrolysis in cells. In conclusion, PhDY-Chol allowed direct visualization of CE homeostasis, validating the function of LIPA in the CE hydrolysis process.

**Fig. 4.**
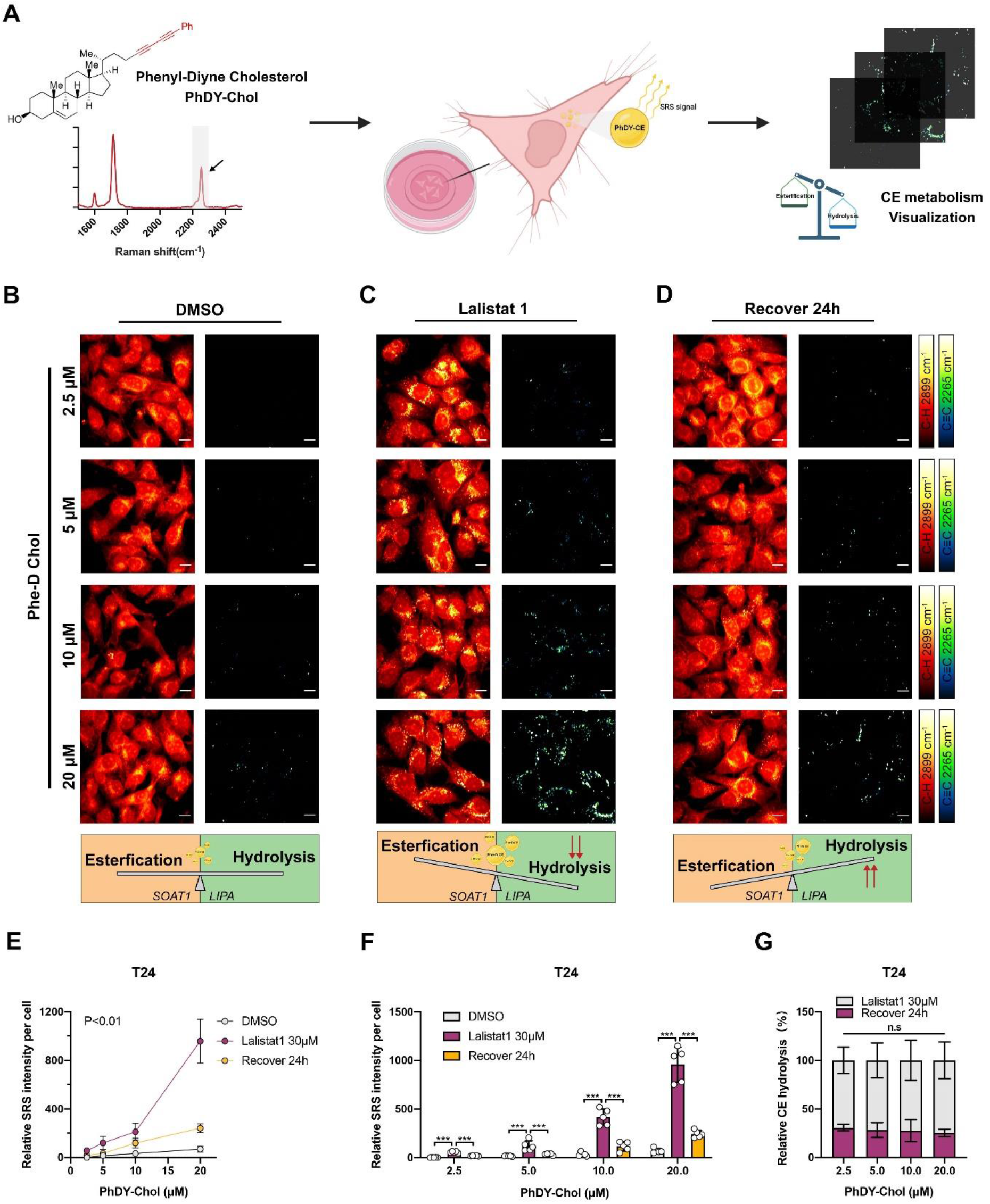
Visualization of CE metabolism by SRS imaging of alkyne-tagged cholesterol. **a** Schematic illustration of Phenyl-Diyne cholesterol visualization by SRS. **b-d** Representative SRS images of T24 after Lalistat 1 treatment and recovery in different concentrations of PhDY-Chol. DMSO (**b**); Lalistat 1 (**c**); Recover 24 h (**d**). Scale bars, 10μm. **e-f** Quantitative analysis of PhDY-Chol in T24 after Lalistat 1 treatment and recovery. Each dot represents a single detection frame. At least 50 cells / sample were imaged. **g** Relative CE hydrolysis percentage of T24 quantitated by using PhDY-Chol. intensity ratio of Lalistat 1 to 24 h recovery. Error bars represent SD of the mean. *** p < 0.001; n.s, not significant.

### LIPA drives rapid hydrolysis of cholesteryl arachidonate

CE is esterified from cholesterol and diverse fatty acids. To profile the fatty acid species involved in LIPA-driven CE hydrolysis, liquid chromatography with tandem mass spectrometry (LC-MS/MS) was employed. Six abundant CEs were then highlighted and analyzed (**Fig. 5a-b**; Supplementary **Fig. 4a-c**). Compared to the control groups, Lalistat 1 dramatically increased the amount of all kinds of CE in both T24 and MDA-MB231 cells, while such increase was mitigated after the recovery. Following the SRS imaging, the fold-changes of both CE accumulation and hydrolysis in T24 were entirely larger than in MDA-MB231 across all CEs. Of concern, there were conspicuous jumps (rise by 160 times in T24, 19 times in MDA-MB231) and slumps (drop by 14 times in T24, 2 times in MDA-MB231) in cholesteryl arachidonate (CE 20:4), demonstrating its high-speed hydrolysis (**Fig. 5a-b**). The relative hydrolysis rate was then defined by the abundance ratio of the Lalistat 1 treated group to the recovery group (**Fig. 5c-d**). As expected, the fastest hydrolysis rates were located in CE 20:4 (11.5 in T24, 10.0 in MDA-MB231), indicating the dominant efficiency of LIPA in CE 20:4 hydrolysis. As previously mentioned, cholesteryl oleate was defined as the dominant form of CE stored in prostate cancers^3,27^. We identified cholesteryl oleate (CE 18:1) as the most abundant CE in both types of cells as well (**Fig. 5e-f**; Supplementary **Fig. 4d-e**), 33.9% in T24 and 59.3% in MDA-MB231. Intriguingly, the percentages of CEs were reassigned after the treatment of Lalistat 1 or the recovery. Although all six CEs showed similar signatures in abundance, the alterations were diverse on the CE percentages. The percentage of CE 20:4 substantially increased by Lalistat 1 treatment (rise from 8.4% to 20.7% in T24, from 11.6% to 24.6% in MDA-MB231) and diminished sequentially by the recovery (fall to 4.1% in T24, 8.2% in MDA-MB231). Such distinctive “rise and fall” hydrolysis pattern was only found in CE 20:4 corresponding to its substantial alternations in abundance (**Fig. 5e-f**; Supplementary **Fig. 4d-e**). While CE 18:1 was the dominant species across different treatments, the representative hydrolysis pattern was not achieved. Collectively, the LC-MS/MS results identified CE 20:4 as the preferred substrate in LIPA-driven hydrolysis.

**Fig. 5.**
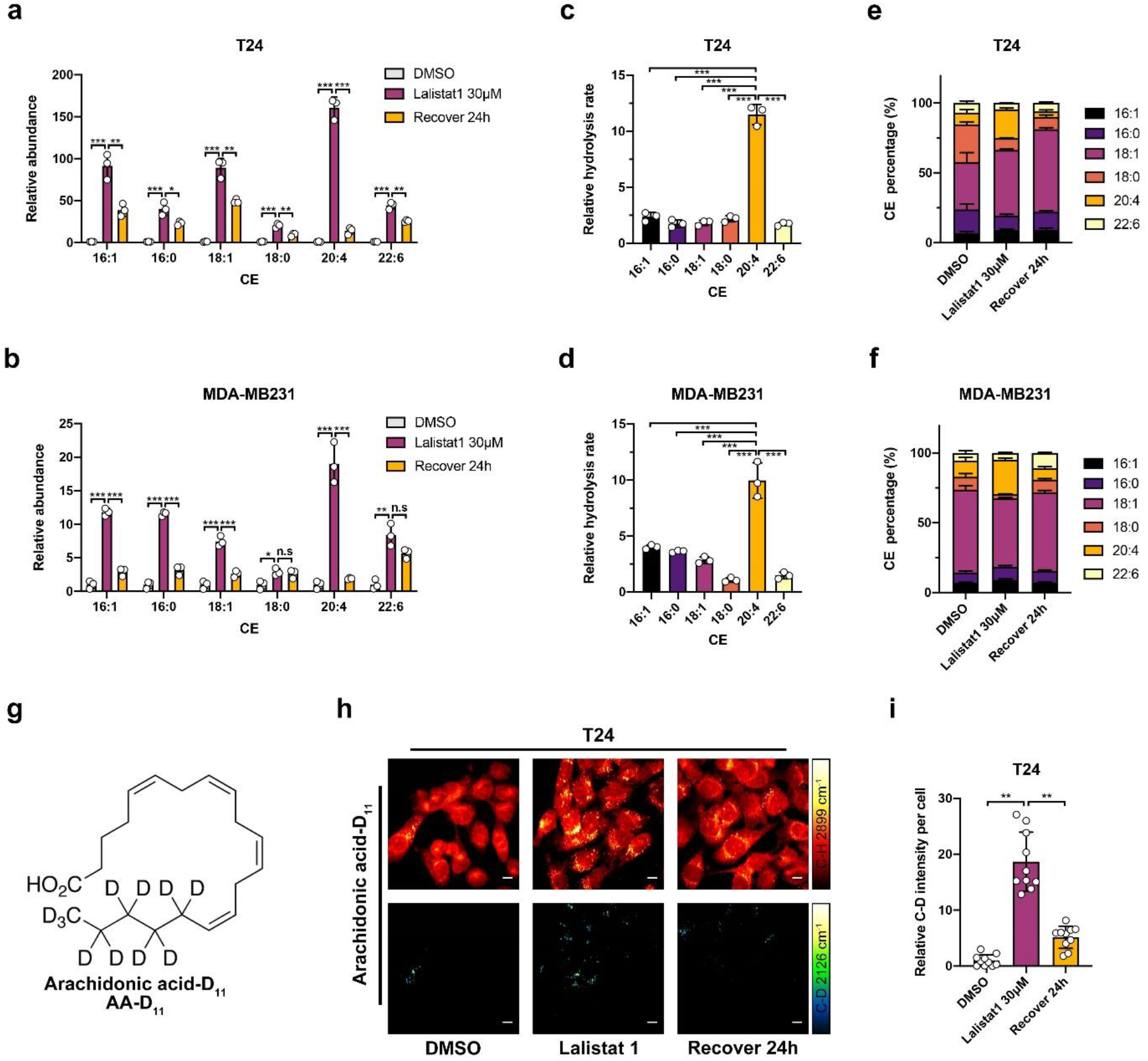
LIPA drives rapid hydrolysis of cholesteryl arachidonate. **a-b** CE profiling by LC-MS/MS after Lalistat 1 treatment and recovery. T24 (**a**); MDA-MB231 (**b**). **c-d** Relative CE hydrolysis rate quantitated by using CE abundance ratio of Lalistat 1 to 24 h recovery. T24 (**c**); MDA-MB231 (**d**). **e-f** Percentage of six CEs after Lalistat 1 treatment and recovery. T24 (**e**); MDA-MB231 (**f**). **g** Schematic chemical structure of AA-D_11_. **h-i** Representative SRS imagines of T24 in C-H and C-D region after Lalistat 1 treatment and recovery(**h**). Scale bars, 10μm. Average C-D SRS intensity in each cell. Each dot represents a single detection frame. At least 100 cells / sample were imaged. Error bars represent SD of the mean. * p < 0.05; ** p < 0.01; *** p < 0.001; n.s, not significant.

Cholesteryl arachidonate derives from arachidonic acid. To validate and visualize the rapid hydrolysis of CE 20:4 driven by LIPA, SRS imaging of deuterated arachidonic acid (arachidonic acid-D_11_, AA-D_11_) was performed (**Fig. 5g**). The C-D stretching vibration from deuterium-labeled arachidonic acid provides the specific 2,126 cm^-1^ peak in the Raman-silent region^28^. By tuning the central beating frequency to be resonant with C-D and C-H stretching vibrations at 2,168 cm^-1^ and 2,899 cm^-1^ respectively, the signals were acquired from AA-D_11_ and C-H biomolecules. Similarly, the excess AA-D_11_ were esterified into LDs. Increased accumulation of AA-D_11_ LDs was found after Lalistat 1 treatment, with corrective reduction after recovery (**Fig. 5h**). During the 24 h recovery, the C-D intensity from AA-D_11_ declined to 28% which was close to the hydrolyzed percentage of PhDY CEs (**Fig. 5i**). The similar hydrolysis rates of PhDY CE and deuterated CE 20:4 indicated the dominant role of CE 20:4 in LIPA-driven hydrolysis. Together, these data show that LIPA drives rapid cholesteryl arachidonate hydrolysis.

### Inhibition of LIPA-driven CE hydrolysis suppresses cancer metastasis

Cholesterol and arachidonic acid, products of cholesteryl arachidonate hydrolysis, are both involved in various cancer phenotypes and contribute to cancer development^2,29^. To address the potential roles of LIPA-driven CE hydrolysis, we performed a series of functional assays by pharmacologic or genetic LIPA inhibition. On basis of the IC_50_ of Lalistat 1, no significant cytotoxicity was found in both T24 and MDA-MB231, even at the high concentration of 120 μM (**Fig. 6a**; Supplementary **Fig. 5a**). Interestingly, enhanced cell viability was found at the high concentration for MDA-MB231 (Supplementary **Fig. 5a**). We next verified the proliferation effect of Lalistat 1 by monitoring two-day viability. Although no statistical significance between Lalistat 1 and the control group (**Fig. 6b**; Supplementary **Fig. 5b**), the average viability of the Lalistat 1 group was higher than the control. Due to the restorable property of Lalistat 1, we asked whether such enhancement results from the rebound of CE hydrolysis. We extended the cell viability assay to the recovery group. After 24 h recovery from Lalistat 1, we found no significant difference in proliferation rate between the two groups (**Fig. 6c**). The long-term effect of Lalistat 1 was next examined by colony formation assay, which showed ineffectiveness as well (**Fig. 6d-e**; Supplementary **Fig. 5c-d**). Whereas of the migration capability, inhibition of LIPA by Lalistat 1 effectively suppressed the migration capacities and partially restored after the recovery (**Fig. 6f-g)**. Such inhibition effects were validated in both MDA-MB231 and MiaPca2 cells (**Fig. 6h**; Supplementary **Fig. 5e**). Although all these three cell lines are defined as mesenchymal cell^30^, their migration capacities were widely varied (Supplementary **Fig. 5f**). Intriguingly, the varied capacities were positively correlated to their CE hydrolysis rate derived from fs-SRS imaging (**Fig. 2d**). To better determine the function of LIPA in cancer proliferation, stable LIPA-knockdown T24 cells (shLIPA) were generated (**Fig. 6i**; Supplementary **Fig. 5g-h**). Similar to the Lalistat 1 treatment, no significant alteration was found in shLIPA T24 cells compared to the control groups in both cell viability and colony formation (**Fig. 6j-k**; Supplementary **Fig. 5i**). Whereas of the migration capability, suppression of LIPA effectively inhibited both cell migration and invasion by 50%, depending on the knockdown efficiency (**Fig. 6l**; Supplementary **Fig. 5j**). These data collectively demonstrate an important role of CE hydrolysis in cancer cell migration.

**Fig. 6.**
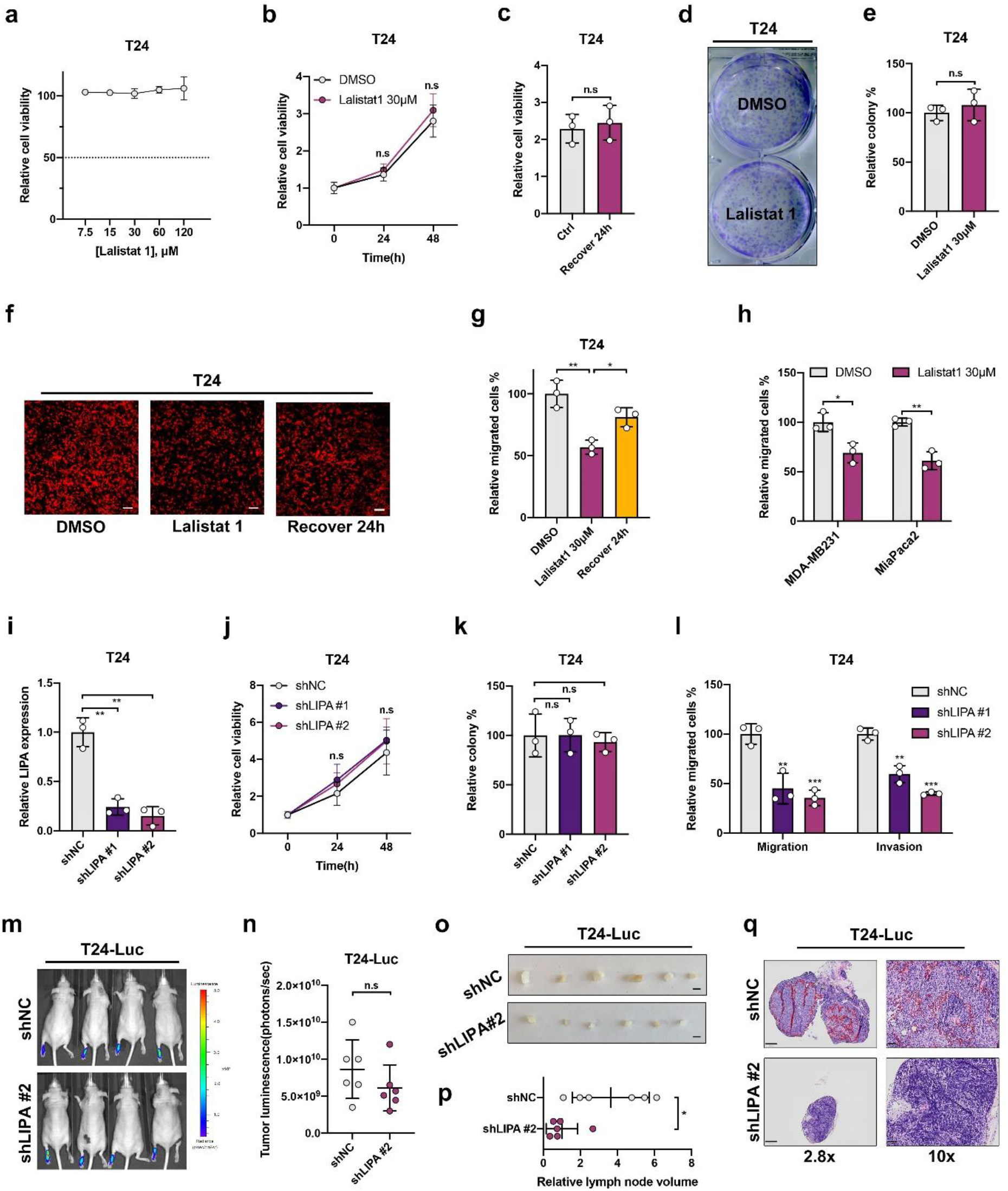
Inhibition of LIPA-driven CE hydrolysis suppresses cancer metastasis. **a** IC50 analysis of Lalistat 1 in T24. **b** Proliferation analysis of T24 after Lalistat 1 treatment. **c** Proliferation analysis of T24 after recovery from Lalistat 1. **d-e** Representative images (**d**) and quantification(**e**) of T24 colony formation after Lalistat 1 treatment. **f-g** Representative images of transwell migration and invasion assay of T24 after Lalistat 1 treatment and recovery (**f**). Scale bars, 100 μm. And quantification of migrated cells(**g**). **h** Quantification of migrated MDA-MB231 and MiaPca2 after Lalistat 1 treatment. **i** RT-qPCR measurement of LIPA mRNA expression level in T24 stably transfected with control shRNA (shNC) or LIPA shRNAs (shLIPA). **j-k** Proliferation (**j**) and colony formation (**k**) analysis of T24-shLIPAs and control. **i** Quantification of transwell migration and invasion by T24-shLIPAs and control. **m-n** Footpad-popliteal lymph node metastasis model by T24-luc. Representative IVIS images of mice injected shLIPA or control cells (**m**) and analysis of tumour luminescence representing xenografts measured after 8 weeks development(**n**). Six mice in each group. **o-p** Representative images of enucleated popliteal lymph nodes (**o**) and quantification of lymph nodes volume (**p**). Scale bars, 2 mm. **q** Representative H&E images of enucleated popliteal lymph nodes. Scale bars, 250 μm and 50 μm. Error bars represent SD of the mean. * p < 0.05; ** p < 0.01; *** p < 0.001; n.s, not significant.

To further explore the function of LIPA-driven CE hydrolysis in cancer progression, we employed a nude mouse model of footpad-popliteal lymph node metastasis. Considering the potential rebound effect of Lalistat 1, only LIPA knockdown cells were used in the mouse model. A luciferase-labeled cell line T24-Luc stably expressing shLIPA and control vector was inoculated into the footpads of nude mice (n = 6 per group). The lymph directly drains from the footpad to the popliteal lymph node, which is an ideal route for cancer metastasis. Thus, the proliferation effect of LIPA is defined by the measurement of primary xenograft in the footpad, while the cancer population in the popliteal lymph node determines the migration capacity. Mice were imaged and harvested after 8-weeks development. The luminescence signals were only detected in the footpad xenografts in both two groups with similar intensities (**Fig. 6m-n**), suggesting the dispensable effect of LIPA during cancer growth in vivo. To evaluate the lymph node metastasis, all the popliteal lymph nodes were harvested from two groups. Strikingly, the volumes of the lymph node in the shLIPA group were smaller than the control group (**Fig. 6o-p**). The H&E stain analyses of the popliteal lymph node were further performed to confirm the metastasis. Consistent with the lymph node size, the areas of lymph nodes were larger invaded with a large number of cancer cells in the control group (**Fig. 6q**). Only a small percentage of infiltrated tumor cells were found in the LIPA knockdown group. Together, our data show that suppression of LIPA-driven CE hydrolysis potently inhibits cancer metastasis, indicating that LIPA can serve as a potential therapeutic target for metastasis.

### A LIPA-CE-NFκB feedback loop

As shown above, CE hydrolysis catalyzed by LIPA effectively contributes to cancer metastasis. To address the interaction network of LIPA during cancer development, cancer cohorts in TCGA were interrogated. The TCGA-BLCA and TCGA-PAAD cohorts were first included to explore the potential pathway correlated with LIPA. Cancer patients were separated into LIPA^High^ and LIPA^Low^ expression groups according to the median expression level of LIPA. The intersections of differential expressed genes (DEGs) were generated (**Fig. 7a**; Supplementary **Fig. 6a**; Supplementary **Table 1**). Of note, the obvious intersection is only found in the upregulated DEGs and next employed in the functional enrichment analysis. The enrichment results confirmed that LIPA is involved in the positive regulation of cell migration (**Fig. 7b**; Supplementary **Table 1**). Several well-known cancer metastasis-related pathways were highlighted, including EMT, NF-κB, PI3K-Akt, and JAK-STAT pathways (**Fig. 7b**; Supplementary **Table 1**). As LIPA-driven CE hydrolysis results in releasing a large amount of arachidonic acid, EMT, NF-κB, and PI3K-Akt were furthered validated due to their verified connection with arachidonic acid^31–33^. The Gene Set Enrichment Analyses (GSEA) of EMT, NF-κB, and PI3K-Akt were performed in four types of cancer cohorts in TCGA. All three pathways showed positive enrichment scores indicating their strong correlations with LIPA-high patients, especially in BLCA (Supplementary **Fig. 6b-e**). As shown in a previous study, inhibition of LIPA impaired Akt phosphorylation in renal cancer^14^. Similarly, we find that both pharmacologic and genetic LIPA repression decreased Akt phosphorylation in T24 (Supplementary **Fig. 6f-g**). Such downregulation was more significant in Lalistat 1 treatment and could be totally restored by the recovery treatment. However, the downregulation of Phospho-S6 Ribosomal protein was only found in Lalistat 1 inhibition. The representative markers in both EMT and NF-κB pathways were then detected. As expected, the expression levels of EMT and NF-κB markers significantly reduced after Lalistat 1 treatment (**Fig. 7c**; Supplementary **Fig.5h**). Of note, the robust rebounds from Lalistat 1 were found in NF-κB target genes, including CXCL8, CSF2, ILB1, IL6, and TNF. Similar inhibition effects were also validated using LIPA knockdown cells (**Fig. 7e**; Supplementary **Fig.5i**). Together, inhibition of LIPA effectively represses the EMT pathway, the PI3K-Akt pathway, and especially the NF-κB pathway.

**Fig. 7.**
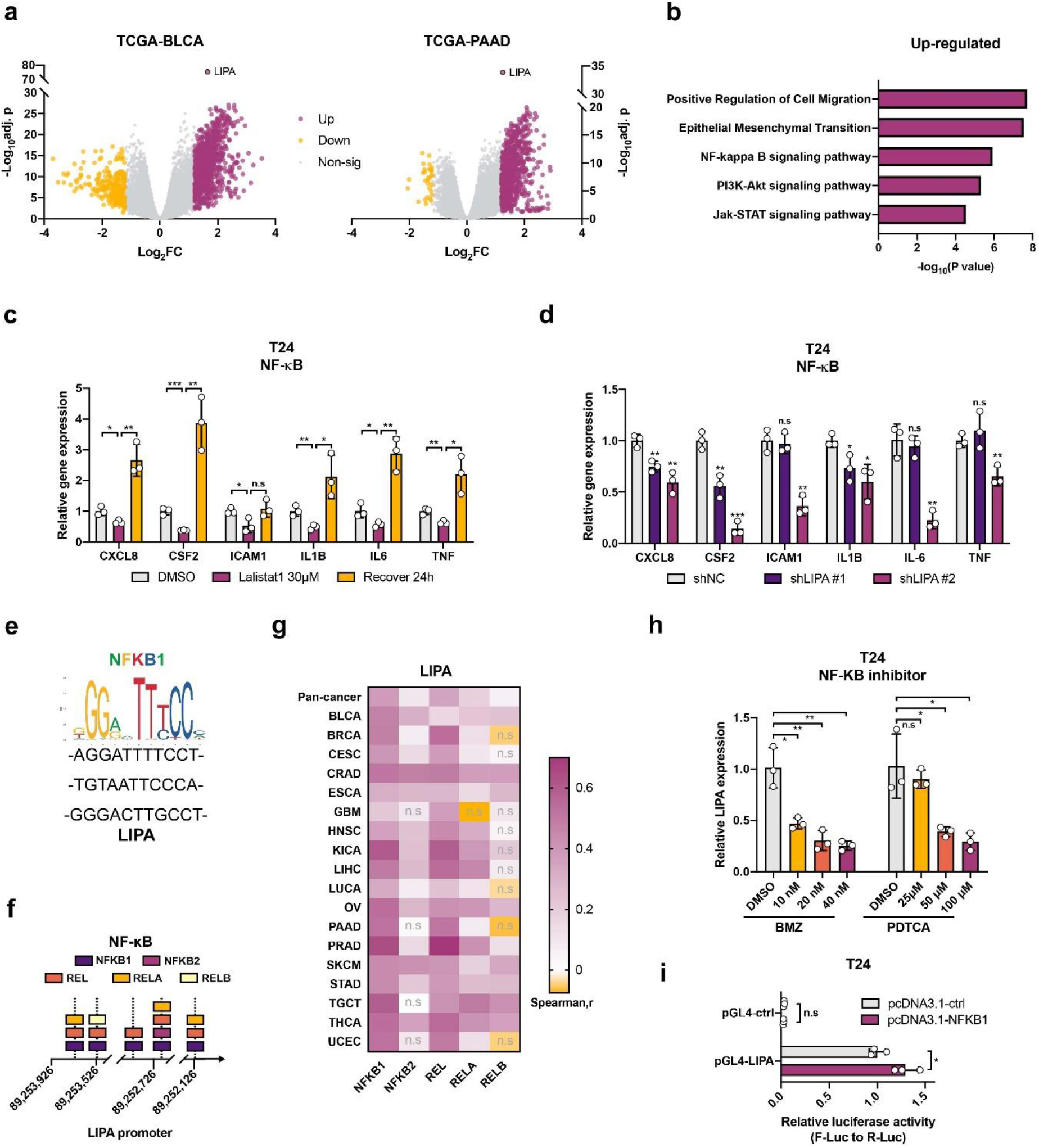
A LIPA-CE-NFκB feedback loop in cancer. **a** The volcano plots of DEGs generated between LIPA^High^ and LIPA^Low^ expression groups in TCGA-BLCA and TCGA-PAAD cohorts. **b** Representative enrichment results for consensus upregulated DEGs. **c-d** RT-qPCR measurement of representative markers involved in NF-κB pathway by pharmacologic (**c**) and genetic (**d**) LIPA inhibition within T24. **e** Conserved motif of NFKB1 and predicted binding sequences in LIPA promoter. **f** Summary of binding sites of NF-κB members in LIPA promoter. **g** A Heatmap of Spearman’s rank correlation coefficient between expression level of LIPA and NF-κB members in various cancer cohorts from TCGA. **h** RT-qPCR measurement of LIPA mRNA expression level in T24 treated with NF-κB inhibitors. **i** Dual-luciferase reporter assay between NFKB1 and LIPA promoter. * p < 0.05; ** p < 0.01; *** p < 0.001; n.s, not significant.

To figure out the upstream regulators of LIPA, in-silico transcription factors perdition analyses were performed on LIPA promoter. Series of transcription factors that potentially bind with LIPA promoter were summarized and sequentially employed in the functional enrichment analysis (Supplementary **Table 2**). Importantly, TNFα signaling via the NF-κB pathway was significantly highlighted across different threshold settings (Supplementary **Fig. 7a-b**; Supplementary **Table 2**). All five NF-κB family members (NFKB1, NFKB2, REL, RELA, RELB) exhibited strong binding specificities with LIPA promoter (**Fig. 7e**; Supplementary **Fig. 7c-f**; Supplementary **Table 2**), especially NFKB1. Of note, most of them were common binding sites of NF-κB members indicating the transcription of LIPA driven by NF-κB dimers (**Fig.7f**; Supplementary **Table 2**). To better understanding the interaction between NF-κB and LIPA, we then detected the expression correlation between LIPA and five NF-κB members by interrogating the TCGA cohorts (**Fig.7g**; Supplementary **Fig. 7g**). Among five members, there were significant positive correlations between LIPA and NFKB1, REL across all cancer type in TCGA (**Fig. 7g**). Cancer patients were next separated into two groups according to NFKB1 and REL expression levels. LIPA expression was significantly higher in the NFKB1^High^REL^High^ patients comparing with the NFKB1^Low^REL^Low^ patients (Supplementary **Fig. 7h-l**). These results collectively suggest that the NF-κB members regulate the level of LIPA. Therefore, the NF-κB pathway inhibitors (Bortezomib, BZM and Pyrrolidinedithiocarbamate ammonium, PDTCA) were applied to verify the potential interaction between the NF-κB pathway and LIPA. Consistent with the above hypothesis, both two NF-κB pathway inhibitors successfully inhibited the expression level of LIPA in a dose dependent manner (**Fig. 7h**). Due to the strong correlation between LIPA and NFKB1, LIPA promoter-driven luciferase reporter and NFKB1 overexpressed vector were further involved to verify the binding potential between LIPA and NFKB1. Compared with the empty vector, the Firefly luciferase activity in NFKB1 overexpressed group significantly increased after normalization with the Renilla luciferase (**Fig. 7i**). These results collectively demonstrate that LIPA expression is positively regulated by NF-κB. Together, CE hydrolysis driven by LIPA promotes the NF-κB pathway, which sequentially enhances LIPA expression, forming a positive LIPA-CE-NFκB feedback loop.

## Discussion

Accumulation of CEs is defined as a functional metabolic hallmark in various cancers^2,34^. However, CE hydrolysis, an essential metabolic step, has been under studied. In this work, we identified LIPA as the dominant regulator of CE hydrolysis in cancers using a combination of bioinformatics, Raman spectroscopy, SRS imaging, and LC-MS/MS. Our results demonstrate that LIPA catalyzes CE hydrolysis, especially the rapid hydrolysis of cholesteryl arachidonate. LIPA-driven hydrolysate promotes cancer metastasis via a NF-κB positive feedback loop (**Fig. 8**). Inhibition of LIPA suppresses cancer metastasis in a mouse model. These results reveal an important role of CE hydrolysis in cancer progression, and open opportunities for treatment of aggressive cancers.

**Fig. 8.**
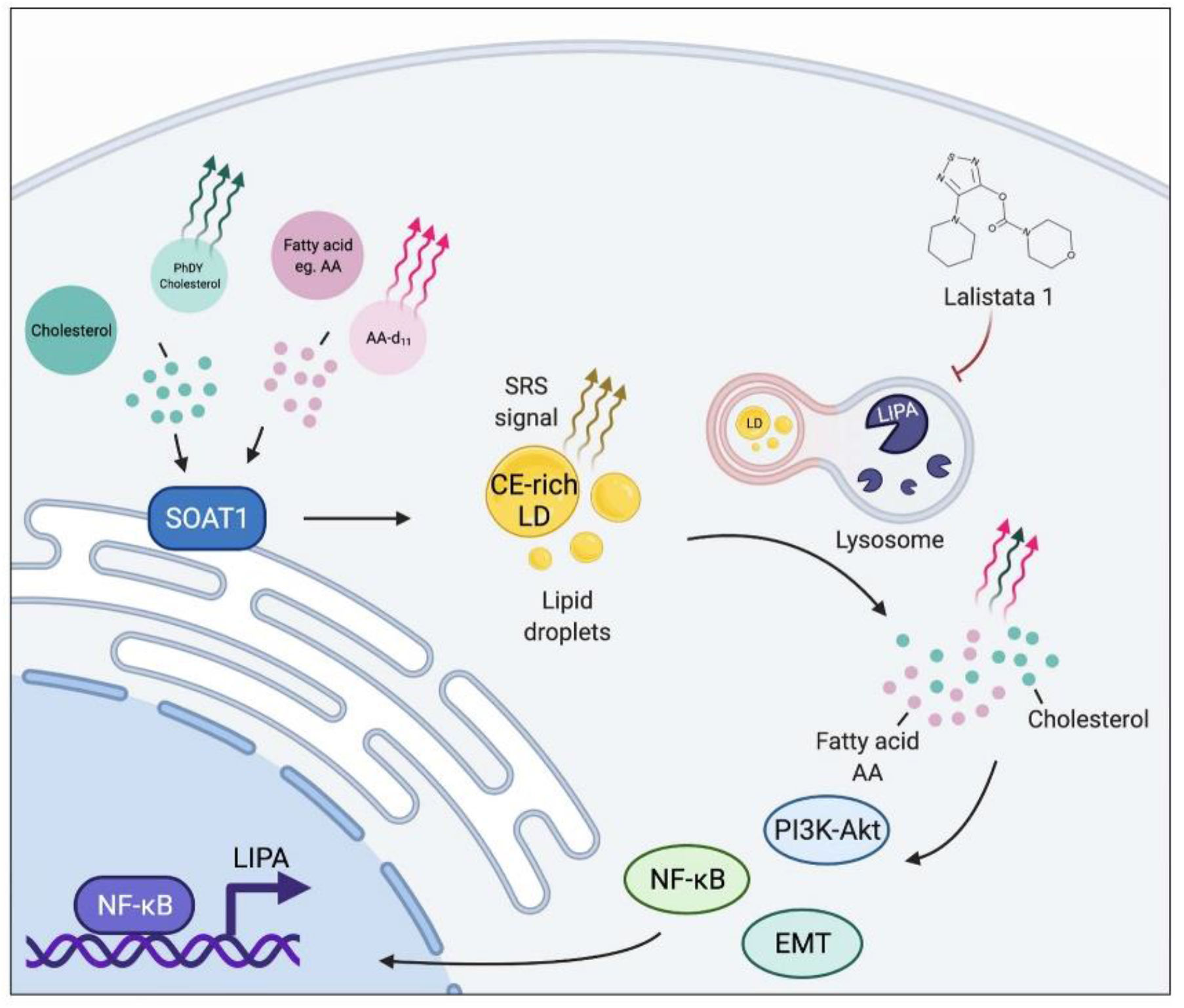
LIPA-driven hydrolysis of cholesteryl arachidonate promotes cancer metastasis by NF-κB positive feedback loop.

LIPA encodes the lysosomal acid lipase (LAL), which functions in the lysosome to catalyze the hydrolysis of CEs and TAGs. LAL deficiency causes Wolman disease and cholesteryl ester storage disease(CESD)^35^. Only a few studies reported the function of LIPA in cancer, including the effect of myeloid-derived suppressor cells^36^ and mesenchymal stem cells^37^ on tumor phenotypes in LIPA knockout mice; up-regulated LIPA in renal cancer promoted tumor proliferation^14^. However, the detailed metabolic characteristics of LIPA in cancer is still underdefined. Herein, CE, instead of TAG, is identified as the major substrate of LIPA. According to CE-targeted LC-MS/MS, only cholesteryl arachidonate(CE 20:4) was further defined as the dominant substrate for LIPA. The robust hydrolysis of CE 20:4 was also directly visualized by SRS chemical imaging of AA-D_11_. Crucially, the recovered hydrolysis rates of PhDY CE and deuterated CE 20:4 were quite similar, demonstrating the prime of CE 20:4 in LIPA-driven hydrolysis. Previously, it has been mentioned that polyunsaturated CE (CE 18:2 and CE 20:4) were preferentially hydrolyzed by LAL in human arterial smooth muscle cells^38^. Of concern, such observation was based on the suppression by chloroquine instead of LIPA-targeted inhibition, while several polyunsaturated CEs were defined as the major substrates. Besides, our study generally characterized the abundance, hydrolysis rate, and CE percentage of six major CEs in cancer cells where the prominent role of CE 20:4 was determined in LIPA-driven hydrolysis. Due to the well-defined function of cholesterol^2^ and arachidonic acid^29^ in cancer, LIPA-driven cholesteryl arachidonate hydrolysis undoubtedly contribute to various cancer phenotypes. Our study highlights that inhibition of LIPA-driven CE hydrolysis suppresses cancer metastasis. Different from previous study in renal cancer^14^, we find no significant alteration of proliferation in both pharmacologic and genetic LIPA inhibition. Such difference might result from the high lipid-dependence in renal cancer cell lines^39^. The complete media were applied in all functional assay where might provide enough exogenous lipid for daily cell growth and minimize the endogenous effect of LIPA. Our data show that the lipid reprogramming, especially the rapid obtainment of free cholesterol and fatty acids from CE hydrolysis, might be critical for cancer metastasis^40^.

Mechanistically, we identified a positive feedback loop involving LIPA, CE hydrolysis, and the NF-kB pathway. Based on the bioinformatics analyses in different TCGA cohorts, we find a significant correlation between LIPA and NF-κB pathway. The NF-κB family has been well defined as the central mediator of the inflammatory process and immune responses. NF-κB activation is mainly driven by inflammatory cytokines and sequentially promotes cancer survival and progression^41,42^. Notably, arachidonic acid does play a vital role in inflammation among many diseased states, including cancer^29,43^. Quite a few studies have highlighted the direct promoting effect of arachidonic acid in NF-κB activation within cancer^32,33,44^. Consistently, our study shows that inhibition of LIPA effectively suppressed NF-κB, especially there were powerful rebounds of NF-κB target genes during the recovery from Lalistat 1. Furthermore, LIPA expression was also positively regulated by NF-κB. Guided by the in-silico binding perdition and expression correlation analyses between NF-κB and LIPA, we verified that NF-κB inhibition decreased LIPA expression and NF-κB bound with LIPA promoter then positively regulated its expression. Notably, the enhancement of LIPA-promoter activity by NFKB1, a member of NF-κB, overexpressed solely was not that prominent suggesting the essential effect of NF-κB dimers in LIPA activation. Interestingly, a potential link between NF-κB and SOAT1 has been established in other models^45,46^. Compared with the acute pharmacologic inhibition of LIPA, the intensity and number of accumulated LDs decreased in the genetic LIPA inhibition by shRNA which might result from the adaptive SOAT1 downregulation from the deactivate NF-κB. Together, NF-κB might function as the central regulator of CE homeostasis. Due to the potential rebound effect of LIPA inhibitor during treatment, NF-κB inhibition could serve as an effective strategy to impair the aberrant CE metabolism in cancer progression. Besides the NF-κB pathway, the EMT and PI3K-Akt pathway were also verified in our study. It is known that such three cancer-promoting pathways have tight interactions^47,48^. Further precise evaluation of LIPA-driven CE hydrolysis in other pathways will help refine the CE metabolic interaction atlas in cancer.

In summary, the current work identified LIPA as the key regulator of CE hydrolysis and determine the LD formation and CE homeostasis in cancer. The rapid hydrolysis of cholesteryl arachidonate driven by LIPA supported the cancer metastasis, intimately coupled with the NF-kB as positive feedback loop. LIPA driven CE hydrolysis represents a vital and targetable metabolic vulnerability in cancer progression, which should be further exploited to fulfill the panorama of cancer metabolism and improve the existing therapeutic strategy.

## Methods

### In silico analysis of CE lipase using online datasets

The pan-cancerous expression analysis and correlation analysis were performed by GEPIA^49^. The overall survival effect was obtained from Human Protein Atlas^50^. All transcriptome datasets for validation obtained from the Cancer Genome Atlas (TCGA) were downloaded using the UCSC Xena browser in Jun 2021. The patients in each cohort were divided into high and low groups based on the median expression level of LIPA. Differentially expressed genes (DEGs) were then generated using the NetworkAnalyst (Supplementary **Table 1**)^51^. All enrichment analyses were performed by the Metascape (Supplementary **Table 1**)^52^. The transcription factor prediction analysis of LIPA promoter was performed by JASPAR (Supplementary **Table 2**)^53^.

### Cell lines and cell culture

Authenticated T24, MDA-MB231, PC-3, and MiaPaca2 cells were obtained from the American Type Culture Collection (ATCC). The T24 cells were cultured in McCoy’s 5A (Gibco, USA), the MDA-MB231 cells were cultured in DMEM (Gibco, USA), the PC-3 cells were cultured in F-12 K (Gibco, USA) and the MiaPaca2 cells were cultured in RPMI 1640 (Gibco, USA). All media were supplemented with 10% fetal bovine serum (Gibco, USA) and 1% penicillin-streptomycin. Cell lines were incubated at 37C° with 5% CO2.

### Chemicals

Lalistat 1, JW480, and arachidonic acid-D11 were purchased from Cayman Chemical. Phenyl-Diyne cholesterol was provided by Dr. Mingji Dai at Purdue University.

### Femtosecond SRS imaging

SRS/pump-probe imaging was performed on a femtosecond SRS microscope. An ultrafast laser system with dual output at 80MHz (InSight DeepSee, Spectra-Physics) provided pump and Stokes beams. The pump and Stikes beams were set to 802 nm and 1045 nm, respectively, to be resonant with the C-H vibration band at 2,899 cm^-1^. For the C-D vibration band at 2,168 cm^-1^, the pump beam was tuned to 852 nm. For the C≡C vibration band at 2,265 cm^-1^, the pump beam was tuned to 846 nm. Stokes beam was modulated by an acoustooptic modulator (AOM, 1205-C, Isomet) at 2.2 MHz. Both beams were linearly polarized. A motorized translation stage was employed to scan the temporal delay between the two beams. Two beams were then sent into a home-built laser-scanning microscope. A 60x water immersion objective lens (NA = 1:2, UPlanApo/IR, Olympus) was used to focus the light into the sample, and an oil condenser (NA = 1:4, UAAC, Olympus) was used to collect the signal. The stimulated Raman loss and pump-probe signals were detected by a photodiode, which was extracted with a digital lock-in amplifier (Zurich Instrument). The power of the pump beam (802nm, 846 nm and 852 nm) and the power of the Stokes beam at the specimen were maintained at ~30mW and 100mW, respectively. The images were acquired at 10 μs pixel dwell time. No cell damage was observed during the imaging procedure.

### Raman spectromicroscopy

Individual LDs in the cells were analyzed by a confocal Raman microscope (LabRAM HR Evolution, Horiba). A 40x water immersion objective (Apo LWD, 1.15 N.A., Nikon) was employed in the system. The sample was excited by a laser of 532 nm. The percentage of CE in the LDs was recorded by measuring Raman intensity at the 702 cm^-1^ peak, which corresponded to cholesterol ester, and the degree of unsaturated fatty acid (UFA) in the cells was recorded by measuring Raman intensity at the 1,654 cm^-1^ peak, which corresponded to C=C stretching vibration, indicating the degree of unsaturated fatty acid. The normalization of Raman intensity was accomplished with the area between 1,400 cm^-1^ to 1,500 cm^-1^, which was the area for -CH2 bending vibration.

### Hyperspectral SRS imaging

Hyperspectral SRS imaging was performed by a spectral focusing approach. The Raman shift is tuned by controlling the temporal delay between two chirped femtosecond pulses. Pump beam and Stokes beam were tuned to 798 nm and 1040 nm to cover the C-H vibration region. Stokes beam was modulated by an acousto-optic modulator (AOM, 1205-C, Isomet) at 2.2MHz. Both beams were chirped by six 12.7cm long SF57 glass rods and then sent to a laser-scanning microscope. A 60x water immersion objective lens (NA =1.2, UPlanApo/IR, Olympus) was used to focus the light on the sample, and an oil condenser (NA = 1.4, U-AAC, Olympus) was used to collect the signal.

To obtain a hyperspectral SRS image, a stack of 100 images at different pump-Stokes temporal delay was recorded. The temporal delay was controlled by an automatic stage, which moved forward with a step size of 10uM, corresponding to ~ 5 cm^-1^. Standard chemicals with known Raman peaks in C-H region from 2,800 to 3,050 cm^-1^, including DMSO, BSA, cholesteryl oleate, and glyceryl trioleate, were used to calibrate the Raman shift to the temporal delay. Hyperspectral SRS images were analyzed using ImageJ. No cell damage was observed during the imaging procedure. Least absolute shrinkage and selection operator (LASSO) unmixing process was done following previously published protocol^26^. This method separates different chemicals in the same window based on their spectral profiles.

### Mass spectrometry analysis

Sample extraction was performed using the Bligh & Dyer protocol^54^. 200μL of ultrapure water was added to each sample. The content was mixed by gentle pipetting followed by 5 minutes of sonication to lyse the cells. The volume of 550μL of methanol (MeOH) and 250μL CHCl3 HPLC grade was added, and the sample was incubated at 4oC for 15 minutes. Next, 250μL ultrapure water and 250μL of CHCl3 HPLC grade were added, resulting in a 2-phase solution. Phase separation was achieved by performing centrifugation at 10,000xg for 5 minutes. The bottom phase (lipids) was transferred to a clean microtube and dried using a speedvac centrifuge. The dried lipid extracts were diluted in 50μL of CHCl3:MeOH 1:1 (v/v) + 10mM ammonium acetate. The volume of 20μL of sample was mixed with 20μL of CE 17:0 300ng/μL. Data was acquired over three minutes for each sample in a triple-quadrupole mass spectrometer (6470 Agilent) equipped with electrospray ion source, operated in the positive ion mode, utilizing precursor ion scan of m/z 369.5. The mass range profiled was m/z 550-800, scan time 3 seconds, collision energy 20V. Ion source parameters were gas temperature 300oC, gas flow 8L/min, nebulizer 20psi and capillary voltage 4kV. An Agilent 1260 HPLC was used to pump CHCl3:MeOH 1:1 (v/v) + 10mM ammonium acetate at 150 μL/min, using flow injection analysis and an injection volume of 15 μL of each sample. The m/z and ion intensity of the ions observed in the mass spectra in were exported to Excel files using the MassHunter Qualitatite Analysis v. B.06.00 for data analysis.

### Cell proliferation and colony formation assays

The cell proliferation assays were performed using MTS reagent (Abcam, UK). All procedures were performed according to the manufacturer’s instructions. For the cell colony formation assay, 2000 cells/well were seeded in a 6-well plate and cultured for 10 days. The colonies were then stained with a 2.5% crystal violet methanol solution for 30 min and scanned to generate images.

### IC_50_ measurement

The IC_50_ measurement for lalistat1 was performed by using 96-well plates. Cells were plated at a density of 5000 cells/well. After 24 hr of seeding, cells were treated with lalistat1 or DMSO from 7.5 μM to 120 μM. Cell numbers were counted after 48 hr of treatment by MTS assay.

### Cell migration and invasion assays

The transwell migration assays were performed by using 24-well transwell chambers with 8 μm pore-sized membranes (Corning, USA). A total of 2×10^5^ cells were plated in the upper chamber with lalistat1 or DMSO treatment. Medium without FBS was added to the upper chamber, and medium with 20% FBS was added to the lower chamber below the cell-permeable membrane. As MDA-MB231 cells and T24 cells have different migration abilities, these two cells need different times for migration. MDA-MB231 cells and MiaPaca2 cells were incubated for 8 hr at 37°C to migrate, and T24 cells were incubated for 6 hr. The transwell membranes were fixed with 10% neutral buffered formalin, and cells that had not migrated through the chamber were removed with a cotton swab. Cells that had migrated through the membrane were stained with 50 μg/mL propidium iodide (Life Technologies, USA) for 30 min and visualized by confocal fluorescence microscopy. The same processes were performed for T24 shLIPA transfected cell lines. Invasion assays were performed using a 6-well invasion chamber. The upper insert contained an 8 μm membrane with a thin layer of MATRIGEL Basement Membrane Matrix (Corning, USA). 1.5×10^6^ cells were plated in the upper chambers. Medium without serum was added to the upper chambers, and medium with 20% FBS was added to the lower chamber. Cells were incubated for 6 hr at 37°C for invasion. The rest processes were similar with migration assay. Images were taken for each well using an FV3000 confocal microscope. The number of migrated/invaded cells was quantified using ImageJ.

### Cell transfection

LIPA shRNA lentivirus was purchased from GenTarget Inc (USA), and the cells were stably infected with non-target control shRNA (shNC) or shRNAs targeting LIPA (shLIPA #1 or shLIPA#2) respectively. The target sequences of shRNAs were listed in Table S3. Both MDA-MB231 and T24 cells were infected with lentivirus at a multiplicity of infection (MOI) of 10. The stable cells were selected using puromycin.

### RNA extraction and quantitative reverse-transcription PCR (RT-qPCR)

Total RNA from cells was extracted using RNeasy Kits (Qiagen, USA) and reverse-transcribed using a iScript™ cDNA Synthesis Kit (Biorad, USA) according to the manufacturer’s instructions. Quantitative PCR was conducted using PowerUp^™^ SYBR^™^ Green Master Mix (Applied Biosystems, USA) and an StepOnePlus^™^ Real-Time PCR System (Applied Biosystems, USA) according to the manufacturer’s instructions. The sequences of the primers used in this study are listed in Table S3. The data analysis was performed using the ΔΔCt method, and expression was normalized to that of RPLP0.

### Mouse model and Fluorescence imaging of Tissue sections

All animal procedures were approved by Boston University IACUC (PROTO201800533). The mouse footpad injection model was used to study the development of metastatic cancer. 6-week-old male NU/J nude mice obtained from the Jackson Laboratory were used. Luciferase-labeled T24 cells were transfected with lentivirus carrying negative control shRNA (T24-Luc shNC) or LIPA targeting shRNA (T24-Luc shLIPA #2). 4×10^6^ cells were inoculated into the footpads of the mice. Lymphatic metastasis was analyzed using a PerkinElmer IVIS Spectrum In Vivo Imaging system. The popliteal lymph nodes were collected after 8-weeks tumor development. Serial sections of popliteal lymph nodes were processed for H&E staining. The tissues were imaged using an Olympus VS120 Automated Slide Scanning Microscope.

### Western blotting

The total cellular protein was extracted using cell lysis buffer (Invitrogen, USA) with 1X protease and phosphatase inhibitor cocktail (Fisher, USA), and 1X DTT (Fisher, USA) and was quantified via a bicinchoninic acid assay (Sigma, USA). The lysates were then separated using SDS–PAGE and electroblotted onto polyvinylidene difluoride membranes. After incubation with the primary and secondary antibodies (Cell Signaling Technology, USA) in sequence, the bands were visualized via enhanced chemiluminescence using an Immobilon Western Kit (Millipore, USA). GAPDH served as a loading control.

### Dual-luciferase reporter assay

The LIPA promoter-Firefly plasmid (pGl4), the NFKB1 overexpressed plasmid (pcDNA3.1), the Renilla plasmid (pRL-TK) and control plasmid were purchased from GenePharma Co (China). Both luciferase activities were measured 48 h later after the transient transfection with different plasmid combinations in accordance with the manufacture’s protocols (Promega, USA). All results were normalized to the Renilla Luciferase activity.

### Statistical analysis

Univariate survival analyses were performed by Kaplan-Meier estimator (Log-rank test). SRS images were quantitatively analyzed using ImageJ software. Raman spectroscopy data were analyzed using Laspec 6 software. LASSO unmixing spectral analysis was performed by MATLAB. Mass spectrometry data were analyzed using Agilent MassHunter Workstation Software Quantitative Analysis. One-way ANOVA or Student’s t test were used for comparisons between groups. Each experiment was performed with a minimum of three biological replicates. All statistical tests were performed using SPSS software version 22.0. Significant differences were considered at * P < 0.05, ** P < 0.01, *** P < 0.001, and n.s means not significant.

## Acknowledgements

This work was supported by R35GM126223, R01CA224275 and R33CA223581 to JXC and R35GM128570 to MD. MD thanks the NIH, Grant P30 CA023168, for supporting shared NMR resources to Purdue Center for Cancer Research. The authors thank Haonan Lin, Xiaowei Ge and Meng Zhang for their help on SRS imaging; thank Yuying Tan for her experimental support; thank Christina Ramires Ferreira from Purdue Metabolomics Facility for their help on mass spectrometry measurements. The research reported in this publication was supported by the Boston University Micro and Nano Imaging Facility and the Office of the Director, National Institutes of Health of the National Institutes of Health under award Number S10OD024993.

## Author contributions

Z.C. and J.X.C. designed and implemented the project. Z.C. and H.J.L. developed the method. Z.C. and F.C. performed experiments. M.D. and X. Y. synthesized Phenyl-Diyne cholesterol. Z.C. and F.C. drafted the manuscript. H.J.L. and J.X.C. reviewed and revised the manuscript.

## Data Availability

The authors declare that the main data supporting the findings of this study are available within the article and its Supplementary Information files. Extra data are available from the corresponding author upon request.

