## Supplementary Figures for "LIPA-driven hydrolysis of cholesteryl arachidonate promotes cancer metastasis via NF-κB"

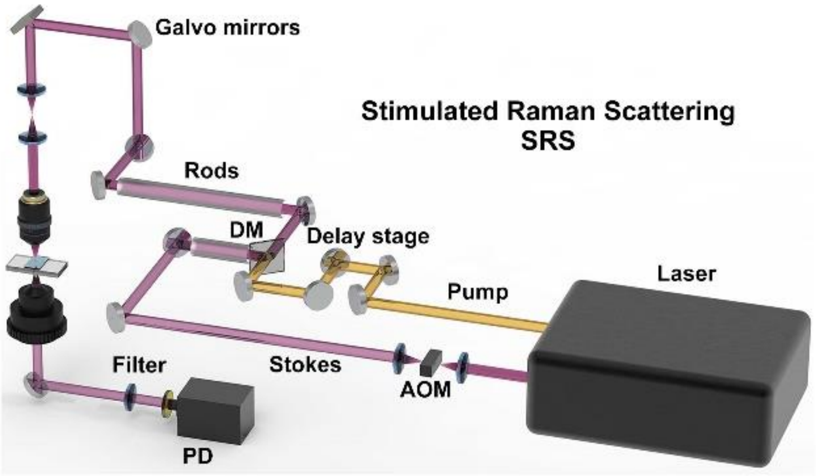


**Supplementary Fig. 1 Schematic illustration of SRS setup**. AOM: acousto-optic modulation. DM: dichroic mirror. PD: photodiode. Rods are removed in fs-SRS.

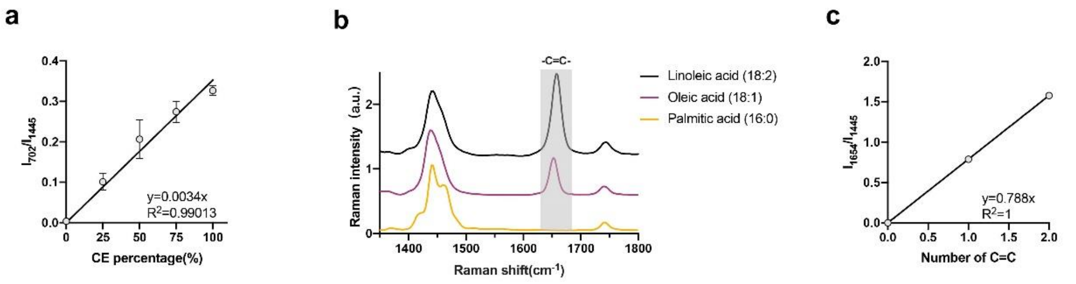


**Supplementary Fig. 2 Quantification of CE percentage and unsaturation degree of intracellular LD. a** Calibration curve of CE percentage in total lipid generated by linear fitting of height ratio between the peak at 702 $cm^{-1}$ ($I_{702}$) and the peak at 1,445 $cm^{-1}$ ($I_{1445}$). Error bars represent SD of the mean. n=3. **b** Raman spectra taken from fatty acids containing different numbers of C=C bonds. **c** Calibration curve of unsaturation degree in total lipid generated by linear fitting of height ratio between the peak at 1,654 $cm^{-1}$ ($I_{1654}$) and the peak at 1,445 $cm^{-1}$ ($I_{1445}$).

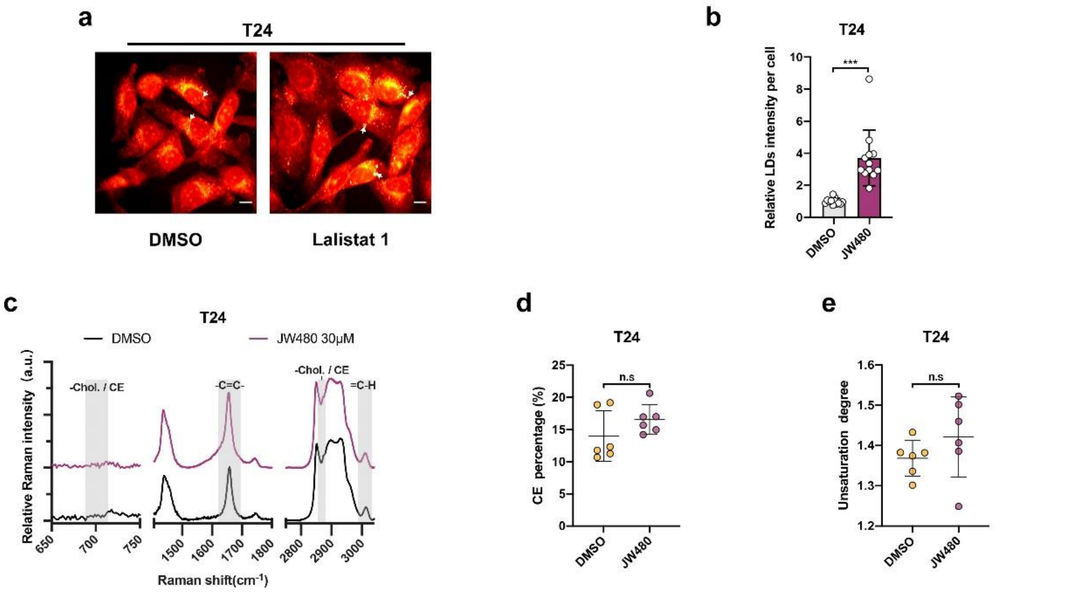


**Supplementary Fig. 3 NCEH1 inhibition by JW480 induces LD accumulation but not CE accumulation in T24 cells. a-b** Representative SRS imagines of T24 after JW480 treatment (**a**). Scale bars, 10μm. Average LDs SRS intensity in each cell (**b**). Each dot represents a single detection frame. At least 100 cells / sample were imaged. **c** Average spontaneous Raman spectra taken from LDs in T24 with JW480 treatment. **d-e** CE percentage (**d**) and unsaturation degree (**e**) in LD of T24 with JW480 treatment. Each dot represents a single cell. At least three LDs / cell were detected. Error bars represent SD of the mean. *** $p < 0.001$; n.s , not significant.

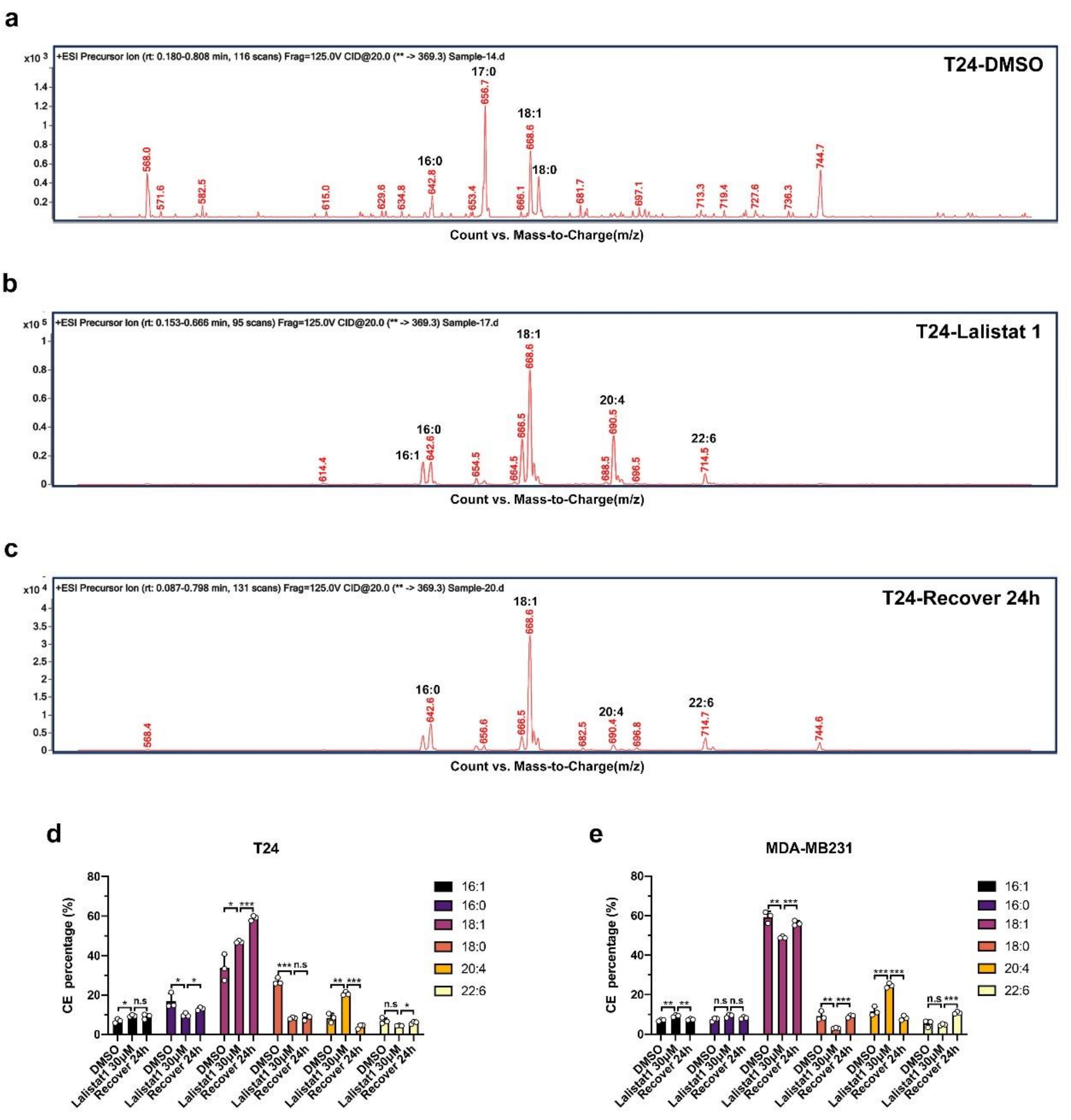


**Supplementary Fig. 4 CE profiling by LC-MS/MS after Lalistat 1 treatment and recovery. a-c** Representative spectra of LC-MS/MS detection in T24 cells treated with DMSO (**a**), Lalistat 1(**b**), and after 24 h recovery (**c**). **d-e** Percentage of six CEs after Lalistat 1 treatment and recovery. T24 (**d**); MDA-MB231 (**e**). Error bars represent SD of the mean.* $p < 0.05$; ** $p < 0.01$; *** $p < 0.001$; n.s , not significant.

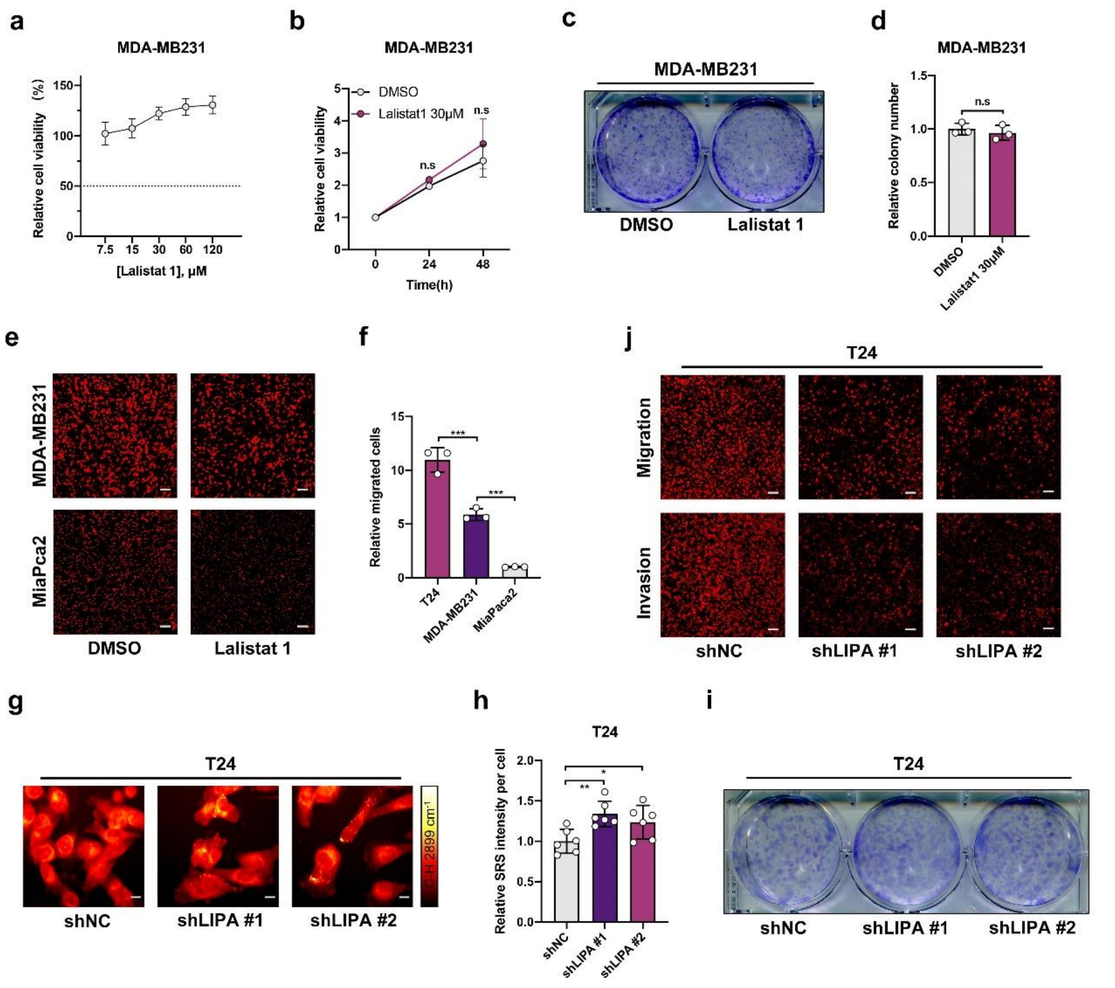


**Supplementary Fig. 5 Inhibition of LIPA-driven CE hydrolysis suppresses cancer invasiveness. a** IC50 analysis of Lalistat 1 in MDA-MB231. **b** Proliferation analysis of MDA-MB231 after Lalistat 1 treatment. **c-d** Representative images (**c**) and quantification(**d**) of MDA-MB231colony formation after Lalistat 1 treatment. **e** Representative images of transwell migration and invasion assay of MDA-MB231 and MiaPca2 after Lalistat 1 treatment. Scale bars, 100 μm. **f** Quantification of migration capability of T24, MDA-MB231, and MiaPca2. **g-h** Representative SRS imagines of T24 stably transfected with control shRNA (shNC) or LIPA shRNAs (shLIPA) (**g**). Scale bars, 10μm. Average LDs SRS intensity in each cell (**h**). Each dot represents a single detection frame. At least 50 cells / sample were imaged. **i** Representative colony formation images of T24-shLIPA and control. Error bars represent SD of the mean.* $p < 0.05$; ** $p < 0.01$; *** $p < 0.001$; n.s , not significant.

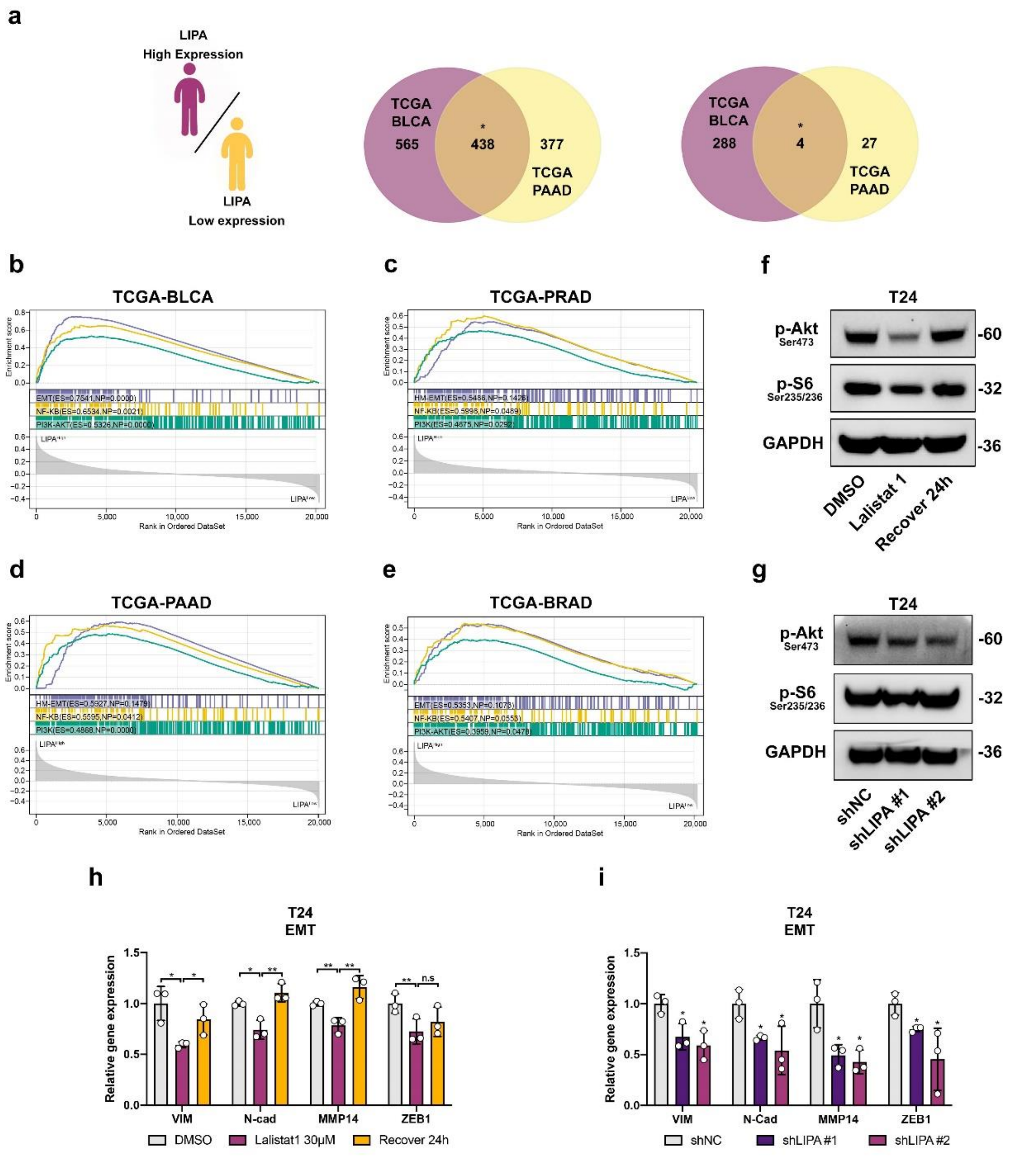


**Supplementary Fig. 6 Inhibition of LIPA suppressed PI3K-Akt and EMT. a** Venn diagrams of consensus DEGs across TCGA-BLCA and TCGA-PAAD cohort. **b-e** GSEA of EMT, NF-κB, and PI3K-Akt performed in four types of TCGA cancer cohorts. **f-g** Western blot of p-Akt and p- S6 by pharmacologic(**f**) and genetic(**g**) LIPA inhibition within T24. **h-i** RT-qPCR measurement of representative markers involved in EMT by pharmacologic(**h**) and genetic(**i**) LIPA inhibition within T24. Error bars represent SD of the mean.* $p < 0.05$; ** $p < 0.01$; n.s , not significant.

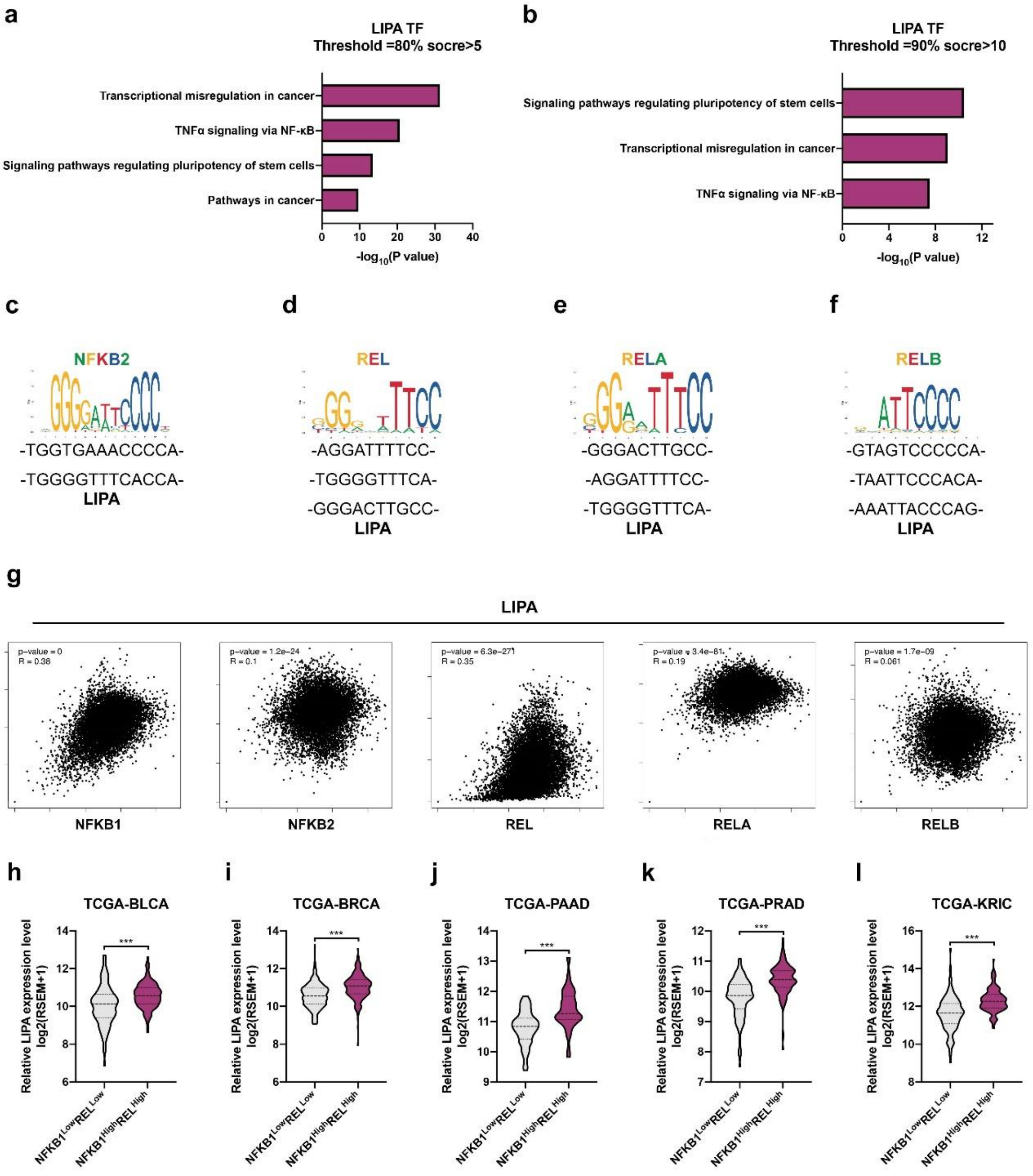


**Supplementary Fig. 7 Binding prediction and expression correlation between NF-κB members and LIPA. a-b** Representative enrichment results for transcription factors that potentially bind with LIPA promoter. **c-f** Conserved motif of NF-κB members and predicted binding sequences in LIPA promoter. NFKB2(**c**); REL(**d**); RELA(**e**); RELB(**f**). **g** Correlation plot between LIPA and NF-κB members in pan-cancer cohorts of TCGA. **h-l** LIPA expression was significantly higher in the NFKB1$^{High}$REL$^{High}$ patients comparing with the NFKB1$^{Low}$REL$^{Low}$ patients in different TCGA cohorts.
