## Supplementary Table 3 for "LIPA-driven hydrolysis of cholesteryl arachidonate promotes cancer metastasis via NF-κB"

| qPCR | | |
| --- | --- | --- |
| **Genes** | **Forward primer** | **Reverse primer** |
| LIPA | ACCAGAGTTATCCTCCCACATAC | AGTCAAGATGCTCCCATTCC |
| VIM | GTGAATACCAAGACCTGCTC | GTTTCGTTGATAACCTGTCCA |
| N-cad | AGAAGACCAGGACTATGACTTGAG | CACCACTACTTGAGGAATTAAGGG |
| MMP14 | TGCCTACCGACAAGATTGATG | ATCCCTTCCCAGACTTTGATG |
| ZEB1 | TAAGCAAACGATTCTGATTCCC | CCTCTACATTTGATACTCCTTCTG |
| CXCL8 | ACTGAGAGTGATTGAGAGTGGAC | AACCCTCTGCACCCAGTTTTC |
| CSF2 | TCCTGAACCTGAGTAGAGACAC | TGCTGCTTGTAGTGGCTGG |
| ICAM1 | TTGGGCATAGAGACCCCGTT | GCACATTGCTCAGTTCATACACC |
| IL1B | TTCGACACATGGGATAACGAGG | TTTTTGCTGTGAGTCCCGGAG |
| IL6 | ACTCACCTCTTCAGAACGAATTG | CCATCTTTGGAAGGTTCAGGTTG |
| TNF | GAGGCCAAGCCCTGGTATG | CGGGCCGATTGATCTCAGC |
| RPLP0 | GAAACTCTGCATTCTCGCTTC | GGTGTAATCCGTCTCCACAG |

Supplementary Table 3. Characteristics of the corresponding shRNAs and qPCR primers included in the context.

| **shRNA** | **Sequence** |
| --- | --- |
| shLIPA #1, TRCN0000029245 | GCCAGGCTGTTAAATTCCAAA |
| shLIPA #2, TRCN0000029247 | GCAGATGTCTACGACGTCAAT |
| shNC | GTCTCCACGCGCAGTACATTT |
